# Deep learning based *k*_cat_ prediction enables improved enzyme constrained model reconstruction

**DOI:** 10.1101/2021.08.06.455417

**Authors:** Feiran Li, Le Yuan, Hongzhong Lu, Gang Li, Yu Chen, Martin K. M. Engqvist, Eduard J Kerkhoven, Jens Nielsen

## Abstract

Enzyme turnover numbers (*k*_cat_ values) are key parameters to understand cell metabolism, proteome allocation and physiological diversity, but experimentally measured *k*_cat_ data are sparse and noisy. Here we provide a deep learning approach to predict *k*_cat_ values for metabolic enzymes in a high-throughput manner with the input of substrate structures and protein sequences. Our approach can capture *k*_cat_ changes for mutated enzymes and identify amino acid residues with great impact on *k*_cat_ values. Furthermore, we applied the approach to predict genome scale *k*_cat_ values for over 300 yeast species, demonstrating that the predicted *k*_cat_ values are consistent with current evolutional understanding. Additionally, we designed an automatic pipeline using the predicted *k*_cat_ values to parameterize enzyme-constrained genome scale metabolic models (ecGEMs) facilitated by a Bayesian approach, which outperformed the default ecGEMs in predicting phenotypes and proteomes and enabled to explain phenotype differences among yeast species. The deep learning *k*_cat_ prediction approach and automatic ecGEM construction pipeline would thus be a valuable tool to uncover the global trend of enzyme kinetics and physiological diversity, and to further elucidate cell metabolism on a large scale.

## Introduction

Enzyme turnover number (*k*_cat_), which defines the maximum chemical conversion rate of a reaction, is a critical parameter for understanding metabolism, proteome allocation, growth and physiology of a certain organism^1–3^. There are large collections of *k*_cat_ values available in the enzyme databases BRENDA^4^ and SABIO-RK^5^, which are, however, still scarce compared to the variety of existing organisms and metabolic enzymes, largely due to the lack of high-throughput methods for *k*_cat_ measurements. Additionally, the experimentally measured *k*_cat_ values have considerable variabilities due to varying assay conditions such as pH, cofactor availability and experimental methods^6^. Altogether, the sparse collection and considerable noise limit the usage of *k*_cat_ data for global analysis and may mask the enzyme evolution trend.

In particular, enzyme-constrained genome scale metabolic models (ecGEMs), where the whole-cell metabolic network is constrained by enzyme catalytic capacities and thereby able to accurately simulate maximum growth ability, metabolic shifts and proteome allocations, rely heavily on genome scale *k*_cat_ values^2,7^. Even for well-studied organisms, the *k*_cat_ coverage is far less than complete^8–10^. When data are missing, ecGEMs usually use assumed *k*_cat_ values from similar reactions or adopt available *k*_cat_ values from other organisms, which could cause model predictions deviating from experimental observations^7^. Thus, there is a clear requirement for obtaining a large scale of *k*_cat_ values to improve the model accuracy and get more reliable simulations for delicate phenotypes^11^.

Previously, machine learning has been used to predict *k*_cat_ values based on features such as average metabolic flux and the catalytic sites obtained from protein structures^9^. Due to the requirement of feature data and absolute proteome data in the training dataset, this approach was only applied to the most well-studied bacterium *Escherichia coli*, thus limiting its usage for large scale prediction of *k*_cat_ values for multiple organisms. In contrast, deep learning does not rely on feature selection and has been applied and shown great performance in modeling chemical space^12^, gene expression^13^, enzyme related parameters such as enzyme affinity^14^, and enzyme commission numbers (EC numbers)^15^.

Inspired by these efforts, we developed a deep learning model and demonstrated its capability for large scale prediction of *k*_cat_ values, as well as for identifying key amino acid residues that affect these predictions. We showcased the predictive power of the deep learning model by predicting genome scale *k*_cat_ profiles for 343 yeast/fungi species, accounting for more than 300,000 enzymes and 3,000 substrates. The predicted *k*_cat_ profiles enabled reconstruction of 343 ecGEMs for the yeast/fungi species through an automatic Bayesian based pipeline, which can accurately simulate growth phenotype among yeast species and identify the phenotype related key enzymes.

## Results

### Construction of a deep learning framework for *k*_cat_ prediction

A deep learning framework was developed by combining a graph neural network (GNN) for substrates and a convolutional neural network (CNN) for proteins (Fig. 1). In this framework, substrates were represented as molecular graphs converted from SMILES (the simplified molecular-input line-entry system) and protein sequences were split into overlapping n-gram amino acids. To train the neural network, we generated a comprehensive dataset from the BRENDA^4^ and the SABIO-RK database^5^. Several rounds of data preprocessing and cleaning were performed to filter out incomplete entries with missing information and redundant entries across databases, to ensure that the dataset contains unique entries with substrate name, substrate SMILES, EC number, protein sequence, organism name and *k*_cat_ value information. The final dataset contained 16,838 unique entries catalyzed by 7,822 unique protein sequences from 851 organisms and converting 2,672 unique substrates (Supplementary Figure 1-2). This dataset was randomly split into training, validation and test dataset by 80%, 10%, and 10%, respectively.

**Figure 1.**
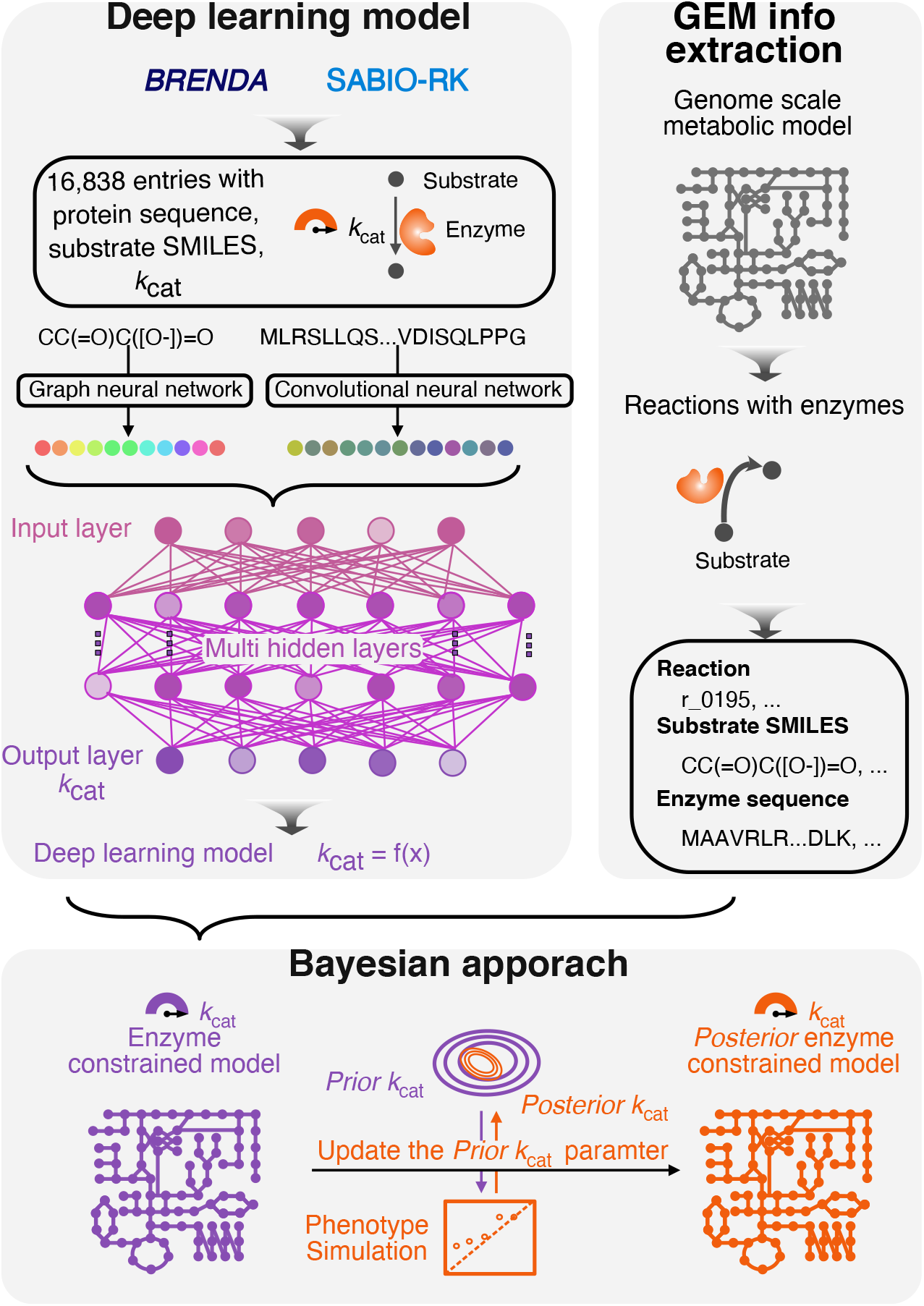
Deep learning of enzyme turnover numbers (*k*_cat_) for genome scale metabolic model (GEM) parameterization. Firstly, we developed an approach for *k*_cat_ prediction by combining a graph neural network (GNN) for substrates and a convolutional neural network (CNN) for proteins. Secondly, we extracted information from GEMs as the input for the deep learning model to predict *k*_cat_ values. Thirdly, we developed a Bayesian facilitated pipeline to reconstruct enzyme-constrained GEMs (ecGEMs) using the predicted *k*_cat_ profiles from deep learning model.

### Deep learning model performance for *k*_cat_ prediction

We first evaluated the effects of different model hyperparameters on deep learning performance using learning curves (Supplementary Figure 3). Note that 2-radius subgraphs and 3-gram amino acids used to extract the substrate and protein vectors can considerably improve the deep learning performance compared with other tested hyperparameter settings (Supplementary Figure 3a). When investigating the effect of vector dimensionality, we found that more highly dimensional vectors used for substrates and proteins led to somewhat better performance (Supplementary Figure 3b). Then, Additionally, the model performed much better when the number of time steps/layers in GNN/CNN is 2 or 3 (Supplementary Figure 3c). With the settled parameters (r-radius is 2, n-gram is 3, vector dimensionality is 20, number of time steps in GNN is 3, and number of layers in CNN is 3), the training dataset was used to train the deep learning model. We observed that the Root Mean Square Error (RMSE) of *k*_cat_ prediction in the validation and test datasets gradually decreased with increasing epoch (Fig. 2a), where the number of epochs represents iterations of the dataset passing through the neural network. A final deep learning model was trained and stored for further use, when the RMSE was 0.99 and 1.06 for the validation and test datasets, respectively, signifying that the predicted and measured *k*_cat_ values were overall within one order of magnitude (Fig. 2a). As a result, the deep learning model showed a high predictive accuracy on the original whole dataset and test dataset (Fig. 2b for whole dataset, Pearson’s r = 0.88; Supplementary Figure 4a for test dataset, Pearson’s r = 0.71; Supplementary Figure 4b for test dataset with substrates and enzymes that were not present in the training dataset, Pearson’s r = 0.70). To facilitate the further usage of our deep learning prediction tool, we also supplied a user-friendly example for *k*_cat_ prediction in our GitHub repository with the input of substrate and protein sequence (https://github.com/SysBioChalmers/DLKcat/tree/master/DeeplearningApproach/Code/example).

**Figure 2.**
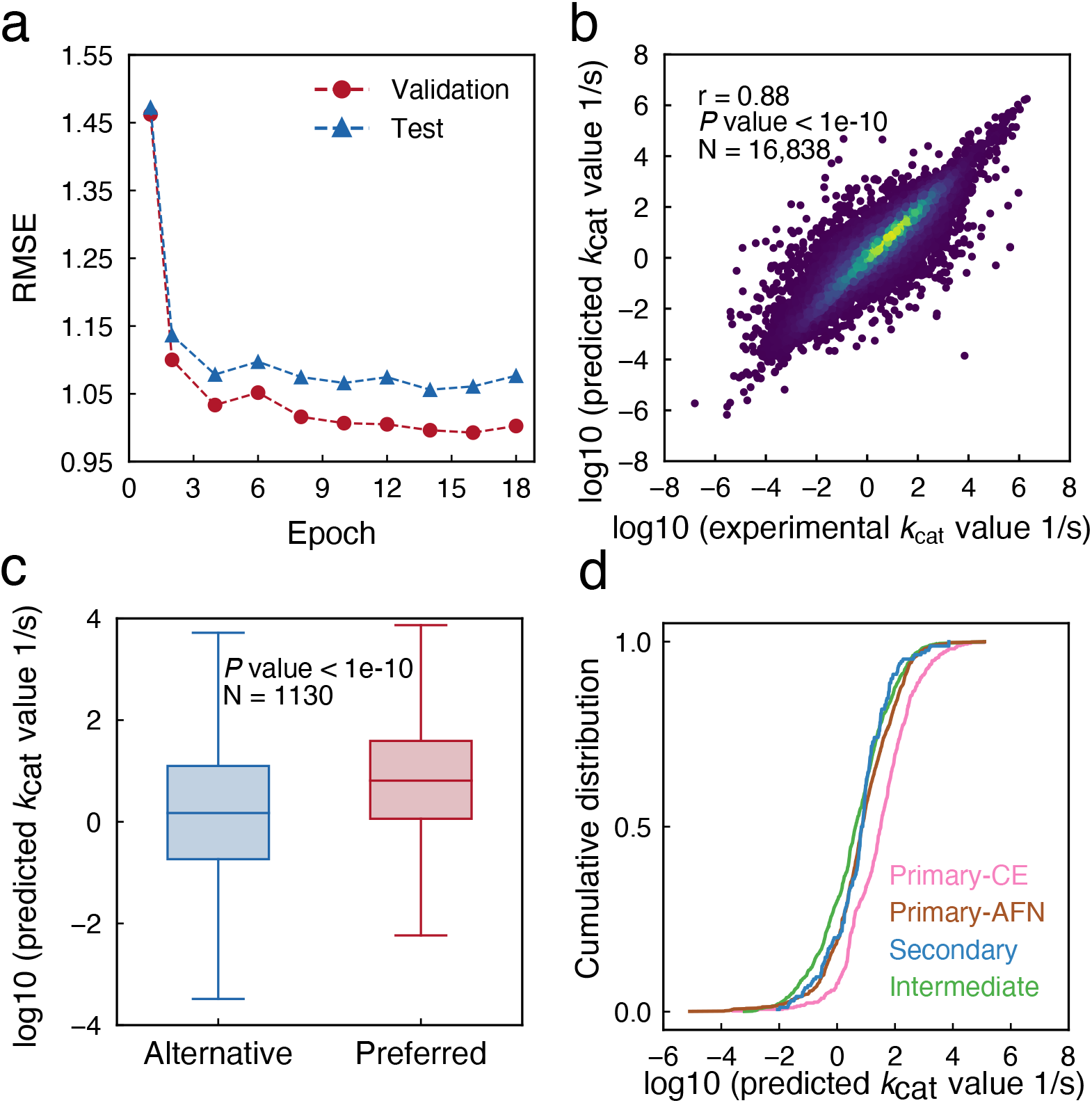
Deep learning model performance for *k*_cat_ prediction. (a) The RMSE of *k*_cat_ prediction during the training process. (b) Performance of the final deep learning model trained by GNN and CNN. The correlation between predicted *k*_cat_ value and those present in the whole dataset was evaluated. The brightness of color represents the density of data points. (c) Enzyme promiscuity analysis on the whole dataset. For enzymes with multiple substrates, we divided the substrates as preferred and alternative by their experimental measured *k*_cat_, then used the predicted *k*_cat_ values for this boxplot. A two-sided Wilcoxon rank sum test was used to calculate *P* value. (d) Cumulative distribution of deep learning-based *k*_cat_ values for enzyme-substrate pairs belonging to different metabolic contexts. Abbreviations: CE, carbohydrate and energy; AFN, amino acids, fatty acids, and nucleotides.

Besides, we investigated whether the deep learning model can identify the preferred substrates for promiscuous enzymes. We classified substrates with the highest *k*_cat_ value for promiscuous enzymes as preferred substrates, and substrates with the lowest one as the alternative substrates, then through comparing the predicted *k*_cat_ values for preferred substrates and alternative substrates (Fig. 2c), we found that our deep learning model are able to predict that the enzymes do indeed have a higher *k*_cat_ for the preferred substrates (median value = 6.45 /s) compared with alternative substrates (median value = 1.49 /s) (*P* value < 1e-10, for promiscuous enzymes in all dataset), which validates the predictive power of our deep learning model in identifying the preferred substrates. The same trend was identified using the prediction for promiscuous enzymes in our test dataset (Supplementary Figure 4c, *P* value = 0.009).

To explore the metabolic contexts for all wildtype enzymes in the original dataset, we mapped these enzymes to four modules on the basis of categorization in KEGG database^16^: primary-CE (enzymes involved in carbohydrate and energy metabolism), primary-AFN (amino acid, fatty acids and nucleotide metabolism), intermediate (metabolism of common biomass components such as cofactors) and secondary metabolism (condition specific metabolism or metabolism related to low concentration metabolites) (Supplementary Table 1). Enzymes associated with primary-CE metabolism on average exhibited a higher predicted *k*_cat_ value than those of primary-AFN, secondary and intermediate metabolism (Fig. 2d), which is in accordance with the previous finding that enzyme-substrate pairs from central carbon metabolism tend to have relatively higher *k*_cat_ values than secondary and intermediate metabolism^6^.

### Prediction and interpretation of *k*_cat_ of mutated enzymes

While the deep learning model displays an overall good performance for predicting *k*_cat_ values (Fig. 2b), we next explored whether the model could capture more details such as the effects of amino acid substitutions on *k*_cat_ values of individual enzymes. To this end, we divided the original annotated dataset into two categories: one including wildtype enzymes and the other mutated enzymes with amino acid substitutions. In these two splits the median *k*_cat_ value of mutant enzymes is lower than that for wildtype enzymes (Supplementary Figure 5a). We found that the deep learning model is a good predictor of *k*_cat_ values for both wildtype enzymes (Fig. 3a for the whole dataset, Pearson’s r = 0.87; Supplementary Figure 5b for the test dataset, Pearson’s r = 0.65) and mutated enzymes (Fig. 3b for the whole dataset, Pearson’s r = 0.90; Supplementary Figure 5c for the test dataset, Pearson’s r = 0.78). Next, several well-studied enzyme-substrate pairs were collected from literature and original dataset from BRENDA^4^ and SABIO-RK^5^ where each enzyme-substrate pair had *k*_cat_ values reported for at least 25 unique amino acid substitutions (Supplementary Table 2). The *k*_cat_ values predicted by the deep learning model correlated very well with the reported experimental *k*_cat_ values (Pearson’s r = 0.94; Fig. 3c). We subsequently divided the entries for each enzyme-substrate pair into two groups based on their experimentally measured *k*_cat_ values: (i) within 0.5-2.0 fold change of the wildtype *k*_cat_ value (‘wildtype-like *k*_cat_’); or (ii) less than 0.5 fold change of the wildtype *k*_cat_ value (‘decreased *k*_cat_’). Scarcity of mutated enzymes with *k*_cat_ values over 2-fold of wildtype *k*_cat_ precluded defining the ‘increased *k*_cat_’ group^17,18^. Using deep learning predicted *k*_cat_ values, we validated that the enzymes from the ‘decreased *k*_cat_’ group indeed showed significantly lower *k*_cat_ values compared to those of enzymes from ‘wildtype-like *k*_cat_’ group for all of the enzyme-substrate pairs (Fig. 3d). The deep learning model is thereby able to capture the effects of small changes in protein sequences on activities of individual enzymes.

**Figure 3.**
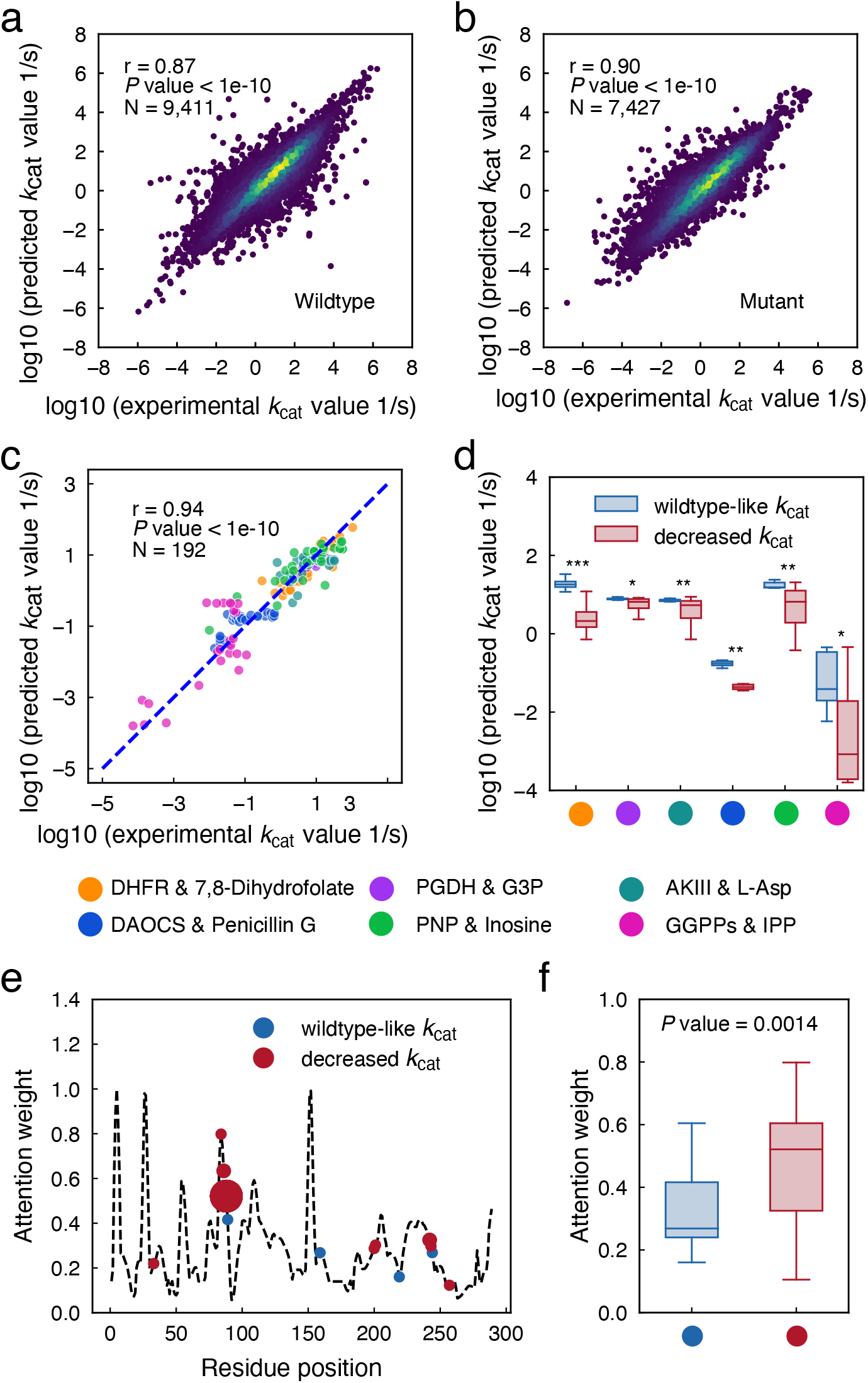
Deep learning model for the prediction and interpretation of *k*_cat_ of mutated enzymes. (a) Prediction performance of *k*_cat_ values for all of the wildtype enzymes via deep learning model. The brightness of color represents the density of data points. (b) Prediction performance of *k*_cat_ values for all of the mutated enzymes via deep learning model. The brightness of color represents the density of data points. (c) Comparison between predicted and measured *k*_cat_ values for several well-studied enzyme-substrate pairs with rich experimental mutagenesis data. Enzyme abbreviations: DHFR, dihydrofolate reductase; PGDH, D-3-phosphoglycerate dehydrogenase; AKIII, aspartokinase III; DAOCS, deacetoxycephalosporin C synthase; PNP, purine nucleoside phosphorylase; GGPPs, geranylgeranyl pyrophosphate synthase. Substrate abbreviations: G3P, glycerate 3-phosphate; L-Asp, L-Aspartate; IPP, isopentenyl diphosphate. (d) Comparison of predicted *k*_cat_ values on several mutated enzyme-substrate pairs between ‘wildtype-like *k*_cat_’ and enzymes with ‘decreased *k*_cat_’. *P* value < 0.05 (*), *P* value < 0.01 (**) and *P* value < 0.001 (***). (e) Attention weight of sequence position in the wildtype PNP enzyme using inosine as the substrate. The mutated enzymes (enzymes with ‘wildtype-like *k*_cat_’ and enzymes with ‘decreased *k*_cat_’) were marked on the curve according to their mutated position. The dot size indicates the number of mutated enzymes occurring in that mutated position. (f) Comparisons of the overall attention weight for the PNP – Inosine pair between enzymes with ‘wildtype-like *k*_cat_’ and enzymes with ‘decreased *k*_cat_’. For two group comparisons in subfigure d and f, a two-sided Wilcoxon rank sum test was used to calculate *P* value.

To investigate which subsequence or amino acid residues dominate enzyme activity, we applied a neural attention mechanism to back-trace important signals from an output of the neural network toward its input^19^. This approach can assign attention weights to each amino acid residue, which then quantitatively describes its importance for the predicted enzyme activity, where higher attention weight signifies higher importance. By this method, we calculated the attention weights for all residues of the *Homo sapiens* enzyme purine nucleoside phosphorylase (PNP) with inosine as substrate, as rich mutation data is available for this enzyme-substrate pair^20^ (Fig. 3e, Supplementary Table 3). Subsequently situating the mutations from the ‘wildtype-like *k*_cat_’ and ‘decreased *k*_cat_’ groups (Fig. 3e) exhibit that mutations from the latter have significantly higher attention weights (Fig. 3f, *P* value = 0.0014, Supplementary Table 4). Mutating amino acid residues with higher attention weights is seemingly having a more substantial effect on enzyme catalytic activity.

### *k*_cat_ prediction for metabolic enzyme-substrate pairs in 343 yeast/fungi species

There are reconstructed GEMs for 332 yeast species plus 11 outgroup fungi^21^, but among these only 14 GEMs were expanded with enzyme-constraints (ecGEMs) due to limited available *k*_cat_ data^2,21^. Thus, we applied the deep learning model to populate enzyme-constrained genome scale metabolic models (ecGEMs). As our developed deep learning model allows prediction of almost all *k*_cat_ values for metabolic enzymes against any substrates for any species except the pair with generic substrates which does not have SMILES information, this enabled generation of ecGEMs for all 343 yeast/fungi species. By using the metabolite and enzyme information extracted from the 343 GEMs^21^ as the input of the deep learning model for *k*_cat_ prediction (Supplementary Figure 6), we predicted *k*_cat_ values for around three million protein-substrate pairs in 343 yeast/fungi species.

By inspecting the global trend for the predicted *k*_cat_ values, we firstly found that yeast and fungal enzymes from primary-CE metabolism have on average the highest *k*_cat_ value compared with enzymes from primary-AFN, secondary and intermediate metabolism (Supplementary Figure 7a), which is consistent with the global trend of all enzymes (Fig. 2c) and literature report^6^. Secondly, we found that specialist enzymes (with narrow substrate specificity) have higher *k*_cat_ values compared with generalist (promiscuous enzymes) that each catalyze more than one reaction in the model (Supplementary Figure 7b). This is aligned with the hypothesis that ancestral enzymes that exhibit broad substrate specificity and low catalytic efficiency improve their *k*_cat_ when they evolve to be a specialist through processes of mutation, gene duplication and horizonal gene transfer. Consistent with reports for *E. coli*^22^, this observation also holds for fungi. Thirdly, we investigated whether sequence conservation trends with *k*_cat_ values. The ratio of non-synonymous over synonymous substitutions, denoted as dN/dS, is commonly used to detect proteins undergoing adaptation^23^. Conserved enzymes with a lower dN/dS have significantly higher *k*_cat_ values compared with relatively lesser conserved enzymes (with high dN/dS), implying that conserved yeast/fungi enzymes under evolutionary pressure are adapted to have higher *k*_cat_ values (Supplementary Figure 7c).

### Bayesian approach for 343 ecGEMs reconstruction

Using the predicted *k*_cat_ values for 343 yeast/fungi species we generated 343 DL-ecGEMs (ecGEMs parameterized with *k*_cat_ values derived from deep learning model prediction). Since the training data for the deep learning model were primarily measured *in vitro*, this implies that also *in vitro k*_cat_ values are predicted by the deep learning model, which is undesired as *in vitro k*_cat_ values can be considerably different from their *in vivo* counterparts^24^. To resolve these uncertainties, we adopted a Bayesian genome scale modeling approach, which has been successfully applied to resolve temperature dependence of yeast metabolism by quantifying and reducing uncertainties in model parameters^25^. Here, we used predicted *k*_cat_ values as mean values for *Prior* distribution and used experimentally measured phenotypes to update it to *Posterior*. The experimental data on yeast/fungi species were collected from literature, collating 445 entries on growth data for 76 species with 16 carbon sources (Supplementary Figure 8, Supplementary Table 5). A sequential Monte Carlo based approximate Bayesian computation (SMC-ABC) approach^25^ was implemented to sample the *k*_cat_ (Methods). The ecGEMs parameterized with the mean values of sampled *Posterior k*_cat_ values were hereafter represented as *Posterior*-mean-DL-ecGEMs.

To test the generality of this SMC-ABC approach and monitor the training process, we first applied this method to ecGEM of *S. cerevisiae*, which has the most abundant experimental data. The experimental phenotype datasets for *S. cerevisiae* were split into training (50%) and test datasets (50%). The training dataset was used to update the *Prior*, which would then be tested on the test dataset after each generation. RMSE between the experimental measurement and prediction for the test dataset was reduced proportionally with the training dataset. After 30 generations, RMSE for the training dataset was 0.5 and for the test dataset was 1, which demonstrates the generalization of the SMC-ABC approach (Supplementary Figure 9).

The Bayesian learning process for *S. cerevisiae* and *Y. lipolytica* are shown as examples (Fig. 4 & Supplementary Figure 10). We calculated RMSE values between measurements and predictions for batch and chemostat growth of *S. cerevisiae* and *Y. lipolytica* under different carbon sources. After several generations, the ecGEMs parameterized with sampled *Posterior k*_cat_ achieved with a RMSE lower than 0.5 (Fig. 4a & Supplementary Figure 10a), which can accurately describe the experimental observations. For instance, the *S. cerevisiae* ecGEM with *Posterior* mean *k*_cat_ values captures the metabolic shift at increasing growth rate (Fig. 4b)—known as the Crabtree effect^26^— while *Y. lipolytica* respires at its maximum growth rate (Supplementary Figure 10b). When exploring which parameters were updated during the Bayesian process, a principal component analysis (PCA) for all 9,500 generated *k*_cat_ sets (95 generations with 100 sets each) showed a gradual move from the *Prior* distribution to the distinct *Posterior* distribution (Fig. 4c for *S. cerevisiae*). The similar gradual move was also observed for *Y. lipolytica* (Supplementary Figure 10c). By comparing the variances of the deep learning and sampled *Posterior k*_cat_ datasets, we found that the Bayesian training process mostly affected variance but not mean predicted *k*_cat_ values (Fig. 4d-e). For *S. cerevisiae*, 2,644 enzyme-substrate pairs reduced their *k*_cat_ variance (Šidák adj. one-tailed F-test *P* value < 0.01), while only 146 pairs changed their mean predicted *k*_cat_ (Šidák adj. Welch’s t test *P* value < 0.01). For the non-conventional yeast *Y. lipolytica*, the value is 2,721 and 159 (Supplementary Figure 10d-e). Consequentially, the sampled *Posterior k*_cat_ has a strong correlation with the deep learning predicted *k*_cat_ (Pearson’s r = 0.83, for *S. cerevisiae*, Fig. 4f; Pearson’s r = 0.83, for *Y. lipolytica*, Supplementary Fig. S10f).

**Figure 4.**
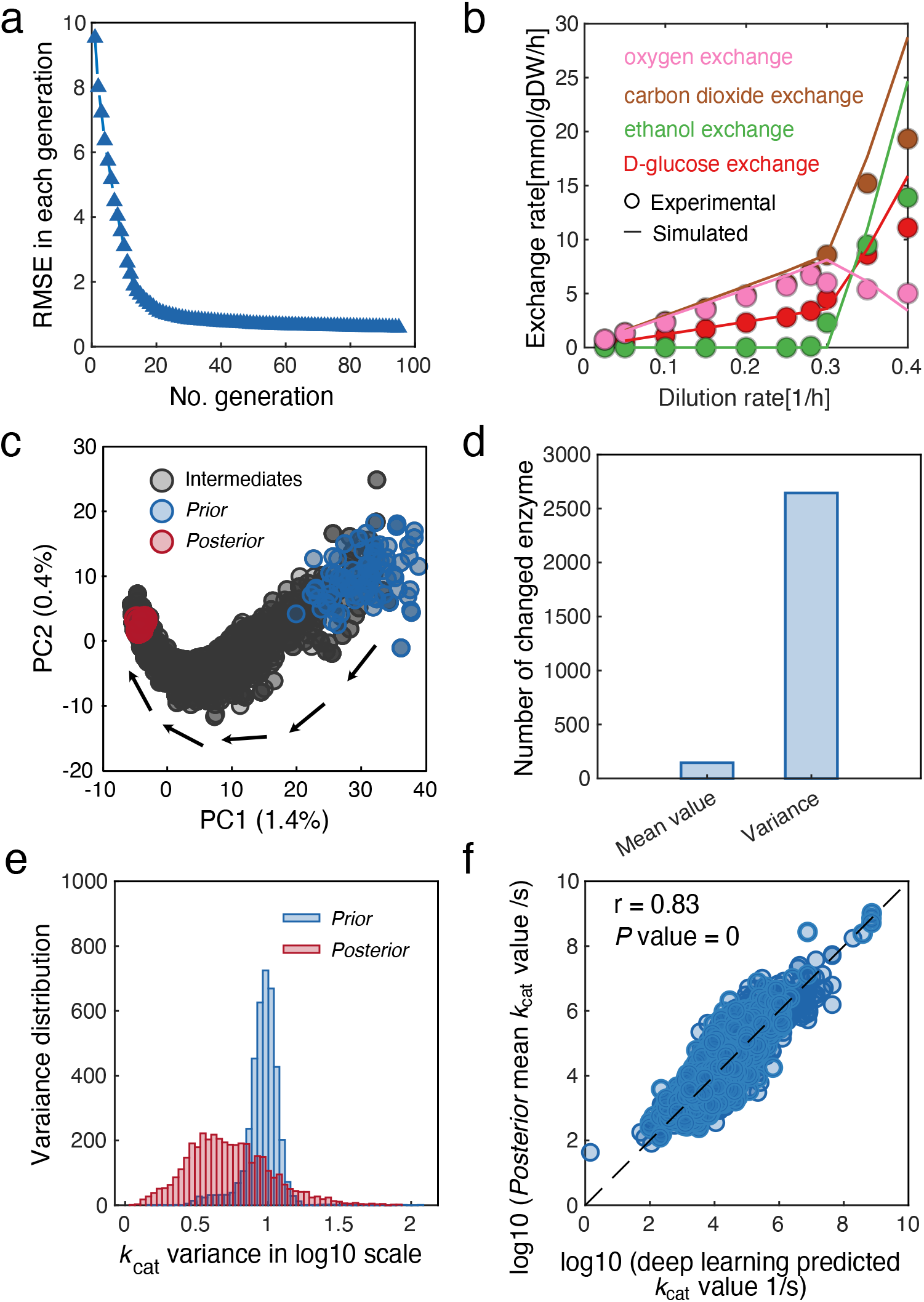
Bayesian modeling training performance for *S. cerevisiae* ecGEM. (a) RMSE for phenotype measurement and prediction during Bayesian training process. (b) Simulated exchange rates by *Posterior*-mean-ecGEM (line) compared with experimental data (dot). *k*_cat_ values in the *Posterior*-mean-ecGEMs here is mean values from 100 sampled *Posterior* datasets after the Bayesian training process. (c) Principal component analysis (PCA) for *k*_cat_ datasets sampled in the Bayesian training approach. Each parameter in the set was standardized by subtracting the mean and then divided by the standard deviation before PCA. Sampled 100 *Prior* datasets are highlighted in blue, while sampled 100 *Posterior* datasets are highlighted in red. All other datasets were termed as “intermediate” and marked in gray. (d) The number of enzymes with a significantly changed mean values (Šidák adjusted Welch’s t test *P* value < 0.01, two-sided) and variance (Šidák adjusted one-tailed F-test *P* value < 0.01) between sampled *Prior* and *Posterior k*_cat_ datasets. (e) Variance distribution comparison for *Prior* and *Posterior* distribution. (f) Correlation between deep learning predicted *k*_cat_ and *Posterior* mean *k*_cat_.

### Deep learning and Bayesian approach improve ecGEMs quality

We subsequently generated *Posterior*-mean-ecGEMs from corresponding DL-ecGEMs for all the 343 yeast/fungi species. For comparison, we also built ecGEMs for the same species with a classical *k*_cat_ parameterization strategy that queried the BRENDA^4^ and SABIO-RK^5^ databases to assign measured *k*_cat_ values to enzyme/reaction pair in the model^2,27^. In case of missing data, certain flexibility was introduced by matching the *k*_cat_ value to other substrates, organisms, or even introducing wild cards in the EC number. This approach is how ecGEMs are routinely parameterized with *k*_cat_ values, and the resulting models are hereafter referred to as Classical-ecGEMs. The Classical-ecGEMs yielded *k*_cat_ values for ca. 40% of enzymes included in the model and generated enzymatic constraints for ca. 60% of the enzyme annotated reactions, while DL-ecGEMs and their derived *Posterior*-mean-ecGEMs covered *k*_cat_ values for ca. 80% of enzymes and defined enzymatic constraints for ca. 90% of enzymatic reactions (Fig. 5a-b). While Classical-ecGEMs have fewer assigned *k*_cat_ values, their reconstruction pipeline also relies heavily on correct enzyme EC number annotations and available measured *k*_cat_ values in the databases, contrasting with the DL-ecGEM reconstruction that relies only on protein sequences and substrate SMILES while resulting in a higher coverage. The missing prediction for DL-ecGEMs and derived *Posterior*-mean-ecGEMs are due to the missing *k*_cat_ prediction for generic substrates which does not have SMILES information.

**Figure 5.**
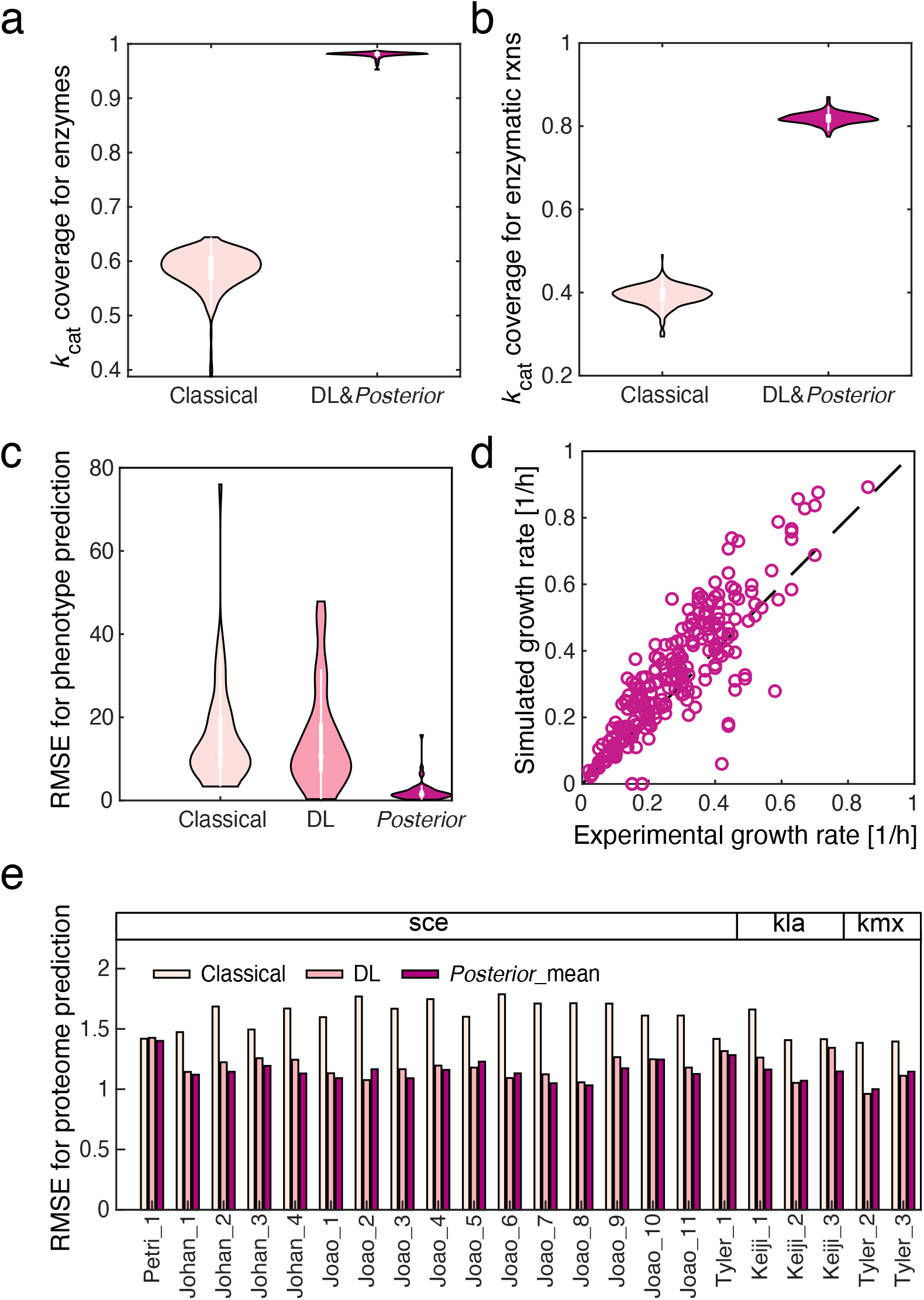
Evaluation of three ecGEM modelling frameworks including Classical-ecGEM, DL-ecGEMs and *Posterior*-mean-ecGEMs. Enzymatic constraint coverage comparison for (a) enzymes and (b) enzymatic reactions. (c) RMSE for the phenotype prediction. (d) Growth prediction for *Posterior*-mean-ecGEMs. (e) Performance of three types of ecGEMs in predicting quantitative proteome data, Classical-ecGEM, DL-ecGEM and *Posterior*-mean-ecGEM are shown. RMSE is shown on log10 scale. Classical-ecGEM is constructed following the pipeline to extract *k*_cat_ profiles from BRENDA and SABIORK, DL-ecGEMs are constructed using the *k*_cat_ profiles predicted from the deep learning model. *Posterior*-mean-ecGEMs here were parameterized by the *k*_cat_ profiles of the mean values from 100 *Posterior* datasets after the Bayesian training process. Detailed conditions for those proteome datasets can be found in the Supplementary Table 6 and collected proteome dataset are available in GitHub repository.

The *Posterior*-mean-ecGEMs and DL-ecGEMs do not only have improved *k*_cat_ coverage but also outperform Classical-ecGEMs in the prediction of exchange rates (Fig. 5c) and are able to predict maximum growth rates in line with the experimentally measured maximum growth rates under different carbon sources and oxygen availabilities (Fig. 5d & more detailed Supplementary Figure 11). Moreover, we used the three types of models to predict required protein abundances and compared this with published quantitative proteomics data from three species with different carbon sources, culture mode and medium setup (Supplementary Table 6). Proteome predictions from *Posterior*-mean-ecGEMs had the lowest RMSE, while DL-ecGEMs already reduced the RMSE by 30% when compared to Classical-ecGEMs (Fig. 5e). Combined, this showed that not only the increased *k*_cat_ coverage but also the Bayesian learning approach contributed to ecGEMs that are better representations of the 343 fungi/yeast species.

### *k*_cat_ profile comparison enables to identify phenotype-related enzyme

The predicted *k*_cat_ values were furthermore able to distinguish between Crabtree positive and negative yeast species. There is much interest in understanding the presence of the Crabtree phenotype among yeast species^28,29^, and a model of *S. cerevisiae* energy metabolism has been used to interpret this phenotype by comparing protein efficiency, i.e. ATP produced per protein mass per time, in its two energy-producing pathways. It was postulated that the Crabtree effect is related to the high yield (HY) pathway (containing Embden–Meyerhof–Parnas (EMP) pathway, tricarboxylic acid (TCA) cycle and electron transport chain (ETC)) having a lower protein efficiency than the low yield (LY) pathway (containing EMP plus ethanol formation) (Fig. 6a)^1^. We here used the *Posterior*-mean-ecGEMs of 102 yeast species (of which 25 are Crabtree positive and 77 are negative with experimental reported phenotype) to similarly calculate protein efficiencies of HY and LY pathways. Of the 102 species we simulated, 89% follow the same trend that Crabtree positive species have a higher LY efficiency while negative species have a higher HY efficiency compared with its LY efficiency, which suggests that Crabtree positive yeast species are more protein efficient using the LY pathway than the HY pathway for producing the same amount of ATP (Supplementary Table 7). For five commonly studied species the results are shown in Fig. 6b, and even though ATP yields in their HY pathways may be different in these species, primarily due to the presence of Complex I, they still follow the same trend (Supplementary Table 7). Inconsistencies in strains where the HY/LY protein efficiency ratio did not trend with the Crabtree effects might be due to additional regulation not considered in ecGEMs^30^.

**Figure 6.**
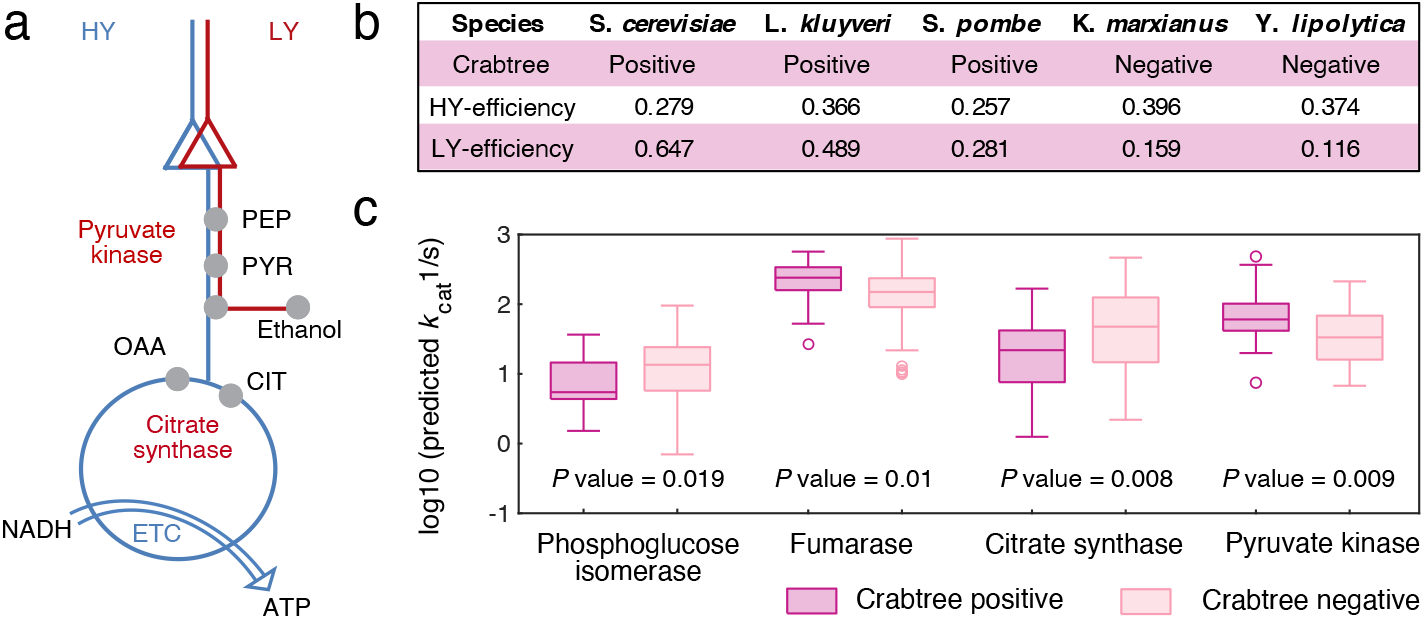
Explanation of the Crabtree effect by energy metabolism. (a) High-yield (HY) and low-yield (LY) pathway definition. (b) Model-inferred protein efficiency of energy metabolism in several common yeast species. Protein efficiency: ATP produced per protein mass per time (Unit: mmolATP/gProtein/h). (c) Enzymes with significantly different *k*_cat_ values between Crabtree positive and negative species. A two-sided Wilcoxon rank sum test was used to calculate *P* value.

With the predicted genome scale *k*_cat_ profiles for yeast species, we can investigate whether key enzymes show significant different *k*_cat_ among 25 Crabtree positive and 77 negative species. Of the enzymes in the energy-producing pathways, only pyruvate kinase, citrate synthase, fumarase and phosphoglucose isomerase had significantly different *k*_cat_ values (Fig. 6c). Since fumarase and phosphoglucose isomerase can operate in reversible direction, it is hard to explain the kinetic effect towards the Crabtree effect. Thus, we would not further discuss the impact of these two enzymes on the Crabtree effect. The *k*_cat_ values of pyruvate kinase were higher in Crabtree positive species compared to negative species (*P* value = 0.009 for deep learning predicted *k*_cat_ values, Fig. 6c). This aligns with a report that increasing pyruvate kinase activity in the Crabtree positive species *Schizosaccharomyces pombe* would increase its fermentation ratio, decrease the growth dependence on respiration and provide resistance to growth inhibiting effects of antimycin A, which inhibits the respiratory complex III^31^. Citrate synthase catalyzes the first and rate-limiting step of the TCA cycle^32^, condensing acetyl-coenzyme A and oxaloacetate to form citrate. We found that the *k*_cat_ of citrate synthase of Crabtree negative species are higher than the Crabtree positive (*P* value = 0.008), which would benefit metabolic flux from entering the TCA cycle (Fig. 6a & 6c). This is consistent with ^13^C-metabolic flux analysis results, which showed that Crabtree negative species have higher TCA flux than Crabtree positive species^33,34^.

## Discussion

The diversity of biochemical reactions and organisms makes it difficult to generate genome scale *k*_cat_ profiles. Here we presented a deep learning model to predict *k*_cat_ values of all metabolic enzymes against all substrates, only requiring substrate SMILES and protein sequences of the enzymes as input, simplifying the feature selection process required for the previous machine learning model^9^. This deep learning approach can therefore be used as a versatile *k*_cat_ prediction tool for any species as long as protein sequence and substrate SMILES are available.

Another advantage of the deep learning model is that it can capture *k*_cat_ changes towards precise single amino acid substitutions. As amino acid substitution is a powerful technique in the enzyme evolution field and is routinely used to probe the enzyme catalytic mechanism^35,36^, it is valuable that attention weight calculation with our deep learning model can identify which amino acid residues have a major impact on the enzyme activity. Particularly, most amino acid substitution experiments performed mutagenesis in the substrate binding site region, since it is hypothesized that the binding region would have a high impact towards the catalytic activity. However, the profound impact remote regions can have towards the catalytic activity has been reported^37,38^. Here, we found high attention weights for the inosine binding region of human PNP enzyme, while also identifying various non-binding residue sites with high attention weight that deserve further validation. In total, our deep learning model is able to predict amino acid substitutions that can impact *k*_cat_ values and thereby serve as part of the protein engineering toolbox^39^.

The deep learning model is able to predict genome scale *k*_cat_ profiles for any species. Phenotype related key enzymes can be identified through comparison of *k*_cat_ values across groups with diverse phenotypes, as done here to identify pyruvate kinase and citrate synthase as Crabtree-effect related enzymes. This approach can as well be applied to identify phenotype related enzymes in other species or even compare among species from different phylogenetic domains. Besides that, global trends in enzyme evolution such as among generalist and specialist enzymes, can be analyzed.

On the other hand, predicted genome scale *k*_cat_ profiles can facilitate the reconstruction of enzyme-constrained models of metabolism. Deep learning predicted *k*_cat_ proved to be a more comprehensive but still practical alternative to matching *in vitro k*_cat_ values from BRENDA^4^ and SABIO-RK^5^ database as is common in Classical-ecGEMs^2,27,40^. Besides the limitation of the EC number annotation for less studied species, *k*_cat_ values measured for the well-studied species are also far away from completeness (Supplementary Figure 1c). For the well-studied species *S. cerevisiae*, only 47 *k*_cat_ values are fully matched with proteins and substrates in the GEM, while other *k*_cat_ values are mostly from fuzzy matching with other substrates, organisms, or even introducing wild cards in the EC number^2^, which also can introduce considerable uncertainty in the reconstructed Classical-ecGEMs. In the earlier published ecGEM reconstruction, a lot of manual work is required to ensure the functionality of Classical-ecGEMs^2^. Compared with the Classical-ecGEM reconstruction, DL-ecGEMs is fully automatic, with reduced uncertainty, significantly increased enzyme coverage and *k*_cat_ coverage for enzymatic reactions and have a more reliable proteome prediction. If there are available experimental growth data, then the ecGEM reconstruction can be further improved through a Bayesian approach. Here, we showed that *Posterior*-mean-ecGEMs are more accurate representatives for their phenotypes and the proteome predictions are also improved, which illustrates how functional ecGEMs can be automatically reconstructed.

In conclusion, we showed how a deep learning approach yields realistic *k*_cat_ which can be used to direct future genetic engineering, understand enzyme evolution, reconstruct ecGEMs that can be used to simulate metabolic flux and phenotype prediction. Besides that, we envision many other possible uses of this deep learning based *k*_cat_ prediction tool such as a novel tool in genome mining and Genome-Wide Association Studies (GWAS) analysis. We also envision this automatic Bayesian ecGEM reconstruction pipeline for further usage in ecGEMs reconstruction, for omics data incorporation and analysis.

## Method and materials

### Preparation of the dataset for deep learning model development

The dataset used for deep learning model construction was extracted from the BRENDA^4^ and SABIO-RK database^5^ on 10 July 2020 by customized scripts via Application Programming Interface (API). We generated a comprehensive dataset including the substrate name, organism information, Enzyme Commission number (EC number), protein ID (UniProt ID), enzyme type, and *k*_cat_ values. Besides, substrate SMILES (Simplified Molecular Input Line Entry System), a string notation to represent the substrate structure, was extracted using substrate name to query the PubChem compound database^41^, which is the largest database of chemical compound information and is easy to access^42^. As different substrates usually have various synonyms in different database and GEMs, we used a customized Python-based script to ensure that the same canonical SMILES could be output for the same substrates with various synonyms, which is essential to help filter redundant entries obtained from different databases (Supplementary Figure 2).

For the BRENDA database^4^, 69,140 entries could be found after downloading and simply processing the accessible data, including 46,417 entries with wildtype enzymes and 22,723 entries with mutated enzymes according to the classification of enzyme type. All these entries contain the required information regarding substrate name, organism, EC number, UniProt ID, enzyme type and *k*_cat_ value. Then we removed duplicates in the entries, and if there are multiple reported measurements for the same enzyme, we only used the maximum value. For the SABIO-RK database^5^, the same data cleaning process was performed. Besides that, we removed the entries with non-standard units for *k*_cat_ values, such as s^(−1)*g^(−1), mol*s^(−1)*g^(−1), J/mol, etc. All *k*_cat_ values were converted to the unit in s^(−1). Available SMILES for substrates were obtained via the API of the PubChem database^41^. Then we combined the dataset extracted from BRENDA database and the SABIO-RK database. Due to high overlap between these two databases, 48,659 unique entries could be found after data cleaning by merging the entries with the same substrate name, EC number, organism, enzyme type and *k*_cat_ value for both databases, and all of the entries have specific substrate SMILES information. Besides the similar approach to keep the maximal values for the multiple measurement, duplicates caused by different synonyms usage in these two databases are filtered using the canonical SMILES. Next, protein sequences are queried with two methods, for entries with UniProt ID information, the amino acid sequences could be obtained via the API of the UniProt database^43^; for entries without UniProt ID, the amino acid sequences were acquired from the UniProt database^43^ and the BRENDA database^4^ based on their EC number and organism information. After that, the sequences of those entries with wildtype enzymes were mapped directly and the sequences of those entries with mutated enzymes were changed according to the mutated sites. Finally, 16,838 entries (including 9,411 entries with wildtype enzymes and 7,427 entries with mutated enzymes) were left as the high-quality dataset for deep learning model construction. Detailed numbers for the data cleaning can be found in Supplementary Figure 2. Data availability: https://github.com/SysBioChalmers/DLKcat/tree/master/DeeplearningApproach/Data/database

### Construction of the deep learning pipeline

In this work, we developed an approach for *in vitro k*_cat_ value prediction by combining a graph neural network (GNN) for substrates and a convolutional neural network (CNN) for proteins. The integration of GNN and CNN can be naturally used to handle pairs of data with different structures, i.e., molecular graphs and protein sequences. In this approach, substrates are represented as molecular graphs where the vertices are atoms, the edges are chemical bonds, and proteins are represented as sequences in which the characters are amino acids.

For substrates, there are just a few types of chemical atoms (e.g., carbon and hydrogen) and chemical bonds (e.g., single bond and double bond). To obtain more learning parameters, we employed r-radius subgraphs to get the vector representations, which are induced by the neighboring vertices and edges within radius r from a vertex^44^. Firstly, substrate SMILES was converted to a molecular graph using RDKit (https://www.rdkit.org). Given a substrate graph, the GNN can update each atom vector and its neighboring atom vectors transformed by the neural network via a non-linear function, e.g., ReLU^45^. Besides, two transitions were developed in the GNN, including vertex transitions and edge transitions. The aim of transitions is to ensure that the local information of vertices and edges is propagated in the graph by iterating the process and summing neighboring embeddings. And the final output of the GNN is a set of real-valued molecular vector representations for substrates.

Similarly, by using the CNN to scan protein sequences, we can obtain low-dimensional vector representations for protein sequences transformed by the neural network via a non-linear function, e.g., ReLU. To apply the CNN to proteins, we defined ‘words’ in protein sequence and split a protein sequence into an overlapping n-gram (n = 1, 2, 3) amino acids^46^. In this work, to avoid low-frequency words in the learning representations, relatively smaller n-gram number of 1, 2 or 3 was set. Also, other important parameters of the neural networks (CNN & GNN) were set as follows: number of layers in CNN: 2, 3 or 4; number of time steps in GNN: 2, 3 or 4; window size: 11 (fixed); r-radius: 0, 1 or 2; vector dimensionality: 5, 10 or 20. These different settings were explored based on R Squared (R^2^) in Equation 1 during the hypermeter tuning to find which hyperparameter is better for improving the deep learning performance. And finally, we used the optimal hyperparameters to train our deep learning model.

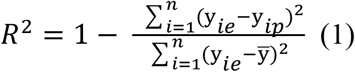

where y*_ip_* is the predicted *k*_cat_ value, y*_ie_* is the experimental *k*_cat_ value, n is the total number of validation dataset.

After the acquisition of the substrate molecular vector representations and the protein sequence vector representations, we concatenated them together and an output vector (*k*_cat_ value) to train the deep learning framework. During the training process, all the datasets were shuffled at the first step, and then were randomly split into training dataset, validation dataset and test dataset at the ratio of 80%:10%:10%. Given a set of substrate-protein pairs and the *k*_cat_ values in the training dataset, the aim of training process is to minimize its loss function. The best model was chosen according to the minimal Root Mean Square Error (RMSE) in Equation 2 on the validation dataset with the least spread between training dataset and validation dataset. For building and training models, the PyTorch v1.4.0 software package was utilized and accessed using the python interface under CUDA/10.1.243.

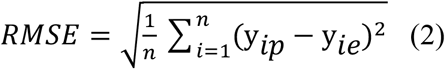

where y*_ip_* is the predicted *k*_cat_ value, y*_ie_* is the experimental *k*_cat_ value, n is the total number of dataset (validation dataset or test dataset).

### Analysis of experimental and deep learning-based *k*_cat_ values across different metabolic contexts

According to the classification of metabolic pathways, metabolic contexts were mainly divided into four different subsystems: primary metabolism-CE (carbohydrate and energy), involving the main carbon and energy metabolism, e.g., glycolysis/gluconeogenesis, TCA cycle, pentose phosphate pathway, etc; primary metabolism-AFN (amino acids, fatty acids, and nucleotides); intermediate metabolism, related to the biosynthesis and degradation of cellular components, such as coenzymes and cofactors; and secondary metabolism, associated with metabolites that are produced in specific cells or tissues, e.g., flavonoid biosynthesis, caffeine metabolism etc^6^. To explore the metabolic subsystems for all of the wildtype enzymes in the experimental dataset, the module in KEGG database^16^ was utilized to assign metabolic pathways for enzyme-substrate pairs by linking the detailed metabolic pathway in KEGG API with EC number annotated in each enzyme-substrate pair. Detailed classification can be found in Supplementary Table 1. Using the trained deep learning model, the predicted *k*_cat_ values were generated for all the enzyme-substrate pairs. The relationship between these predicted *k*_cat_ values and various metabolic contexts was further analyzed, which was compared with the trends of the annotated experimental results.

### Interpretation of the reasoning of deep learning with neural attention mechanism

To interpretate which subsequences or residue sites are more important for the substrate, the neural attention mechanism was employed by assigning attention weights to the subsequences^19^. A higher attention weight of one residue means that residue is more important for the enzyme activity towards the specific substrate. Such attention weights were modeled based on the output of the neural network.

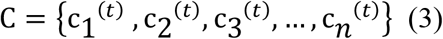

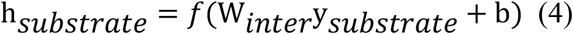

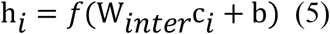

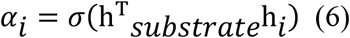

where C is a set of hidden vectors for the protein sequence, c_1_*^(t)^* to c*_n_^(t)^* are the sub-hidden vectors for the split subsequences, y*_substrate_* is the substrate molecular vector, W*_inter_* and b are the weight matrix and the bias vector in the neural network, respectively, f is a non-linear activation function (e.g., ReLU), α*_i_* is the final attention weight value.

For a defined protein, it could be split into overlapping n-gram amino acids and calculated as a set of hidden vectors in Equation 3. Given a substrate molecular vector y*_substrate_* and a set of protein hidden vectors, the substrate embeddings (h*_substrate_*) and subsequence embeddings (h*_i_*) could be output based on the neural network as shown in Equation 4 and Equation 5. By considering the embeddings of y*_substrate_*, the attention weight value for each subsequence was accessible in Equation 6, which represents the importance signals of the protein subsequence towards the enzyme activity for a certain substrate.

### Prediction of *k*_cat_ values for 343 yeast/fungi species

The GEMs of 343 yeast/fungi species were downloaded from the GitHub repository^21^. For each model, all reversible enzymatic reactions were split to forward and backward reactions. Reactions catalyzed by isoenzymes were also split to multiple reactions with one enzyme complex for each reaction. Substrates were extracted from the model and mapped to MetaNetX database to get SMILES structure using corresponding annotated MetaNet IDs for metabolites^47^. Protein IDs for the enzymes were from the model.grRules. Since there are around 200 yeast species are newly sequenced^48^ and are not included in the UniProt database^43^, protein sequences were queried by the protein ID in the protein fasta file for each species (Supplementary Dataset). Reaction IDs, substrate names, substrate SMILES and protein IDs were combined as the input file for the deep learning *k*_cat_ prediction model.

### Analysis of *k*_cat_ values and dN/dS for 343 yeast/fungi species

In a previous study, the genomes of 343 yeast/fungi species combined with comprehensive genome annotations were publicly available^48^. The gene-level dN/dS of gene sequences for pairs of orthologous genes from the 343 species were calculated with yn00 from PAML v4.7^49^. For this computational framework, the input is the single-copy ortholog groups (OGs), and the output is the gene-level dN/dS values extracted from the PAML output files. By mapping the predicted *k*_cat_ values with the gene-level dN/dS values via the bridge of protein ID, a global analysis was performed between the *k*_cat_ values and the dN/dS values for 343 yeast/fungi species across the outgroup (11 fungal species) together with 12 major clades divided by the genus-level phylogeny for 332 yeast species.

### ecGEM reconstruction

ecGEMs are reconstructed by adding enzymatic constraints (Equation 7) into the basic constraints of basic GEMs.

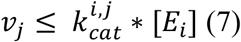

where *v_j_* stands for the metabolic flux (mmol/gDW/h) of the reaction *j*, [*E_i_*] stands for the enzyme concentration for the enzyme *i* that catalyzes reaction *j* and 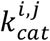 is the catalytic turnover number for the enzyme catalyzing reaction *j*. This constraint is applied to all enzymatic reactions with available *k*_cat_ values.

We used two formats of ecGEMs in the reconstruction process: we adopted the sMOMENT^27^ format in the Bayesian modeling process to speed up the *k*_cat_ mapping process and linear problem construction in the SMC-ABC search; while in the model evaluation and final format, we used the GECKO format to compile all *k*_cat_ values in the model S matrix which would be compatible with all developed GECKO functions^2,50^. There is a developed customized function convertToGeckoModel to facilitate the conversion for these two formats.

Classical-ecGEM reconstruction queries *k*_cat_ values from BRENDA database by matching the EC number, which is heavily relied on the database EC number annotation for the specific species^2,27^. Since more than 200 out of 343 yeast/fungi species are not annotated in UniProt^43^ and KEGG^16^, EC numbers for orthologs annotated in *S. cerevisiae* were borrowed to facilitate Classical-ecGEM reconstruction process for all these 343 species. The *k*_cat_ extraction process used the criteria from the process 13 in the reconstruction methods of the reference^40^.

DL-ecGEM reconstruction extracts all *k*_cat_ values from the deep learning predicted file. To assign *k*_cat_ value for each metabolic reaction, we follow the criteria below 1) *k*_cat_ values predicted for currency metabolites such as H_2_O, H^+^ were excluded; 2) If there are multiple substrates in the reaction, maximum values among substrates were kept; 3) If multiple subunits exist in the enzyme complex, we used the maximum values among all subunits to represent the *k*_cat_ for the complex.

*Posterior*-mean-ecGEM reconstruction uses mean values for accepted *Posterior* distribution. The *k*_cat_ values in the DL-ecGEMs combined with the RMSE (which is 1 in log10 scale) of the *k*_cat_ prediction were used as mean values and variance to make the *Prior* distribution. Each *k*_cat_ value was described with a log normal distribution N(*kcat_i_*, 1). This *Prior* iteratively morphs into a *Posterior* through multiple generations^25^. For each generation, we sampled 128 *k*_cat_ datasets within the distribution, and 100 among those 128 datasets with smaller distance (see next section for the SMC-ABC distance calculation) between phenotype measurements and predictions which can better represent the phenotype were kept to make the distribution for the next generation. Until the distance is lower than the cutoff (RMSE of 0.5), then we accepted the final distribution as *Posterior* distubiton^25^.

### SMC-ABC distance function

Experimental growth data and related exchange rates in batch and chemostat conditions were collected for yeast/fungi species, which are available at Supplementary Table 5. The distance function was designed as RMSE between simulated and experimental values for maximal growth simulations and exchange rates simulations. As for maximal growth simulation, the medium was set in the model by allowing the free uptake of composition, and the objective function was set to maximizing growth. The RMSE was calculated for the simulated and measured growth rates. For the exchange rates simulation, the carbon source uptake rates were constrained based on experimental measurements, and the objective function was also set to maximizing growth. The RMSE was calculated for the simulated and measured exchange rates of all measured exo-metabolites. All measured and simulated rates were normalized by the carbon numbers of the corresponding metabolites before calculation of RMSE. The carbon number for biomass is 41 (mean value for the molecular wight of 1 Cmol biomass of yeast is ∼24.42 g^51^, the biomass equals to 1000 mg). Note that if the substrate or byproduct does not contain any carbon such as O_2_, then the normalizing number is 1. Then the average RMSE of both simulations was used to represent the distance. SMC-ABC search would stop once the RMSE reaches the accepted value or reaches the maximum generation. The accepted value for the distance is set to be lower than 0.5 and the maximum generation is set to be 150.

### Simulations with ecGEMs

We performed different kinds of simulations using the ecGEMs including simulations of growth and protein abundance. Different mediums and growth conditions were set to match the experiment measurement condition, e.g., using xylose as the carbon source or anaerobic condition. Since there are no measured total protein abundance in the biomass for all yeast/fungi species, we used the protein content mass to serve as the total protein abundance for each species and used a sigma factor of 0.5 to serve as the ratio of metabolic protein ratio in total protein abundance.

### Statistical tests for comparison between sampled *Prior* and *Posterior* dataset

Sampled *Prior* and *Posterior k*_cat_ datasets were compared for the difference in the mean values and the variance. Welch’s t test was used to test the significance for the mean values, while one-tailed F-test was used for the reduced variances. The cutoff for the significance was set to 0.01 for the adjusted *P* value corrected by the Šidák method.

### Proteome data collection

All collected proteome data are available in the GitHub repository (https://github.com/SysBioChalmers/DLKcat/tree/master/BayesianApporach/Data/Proteome_ref.xlsx). For relative proteome datasets, we normalized by the identical condition of the absolute proteome data from the literature following the same method as^52,53^. Reference absolute datasets for those relative proteome datasets were documented in the same file.

### Calculation of protein cost and efficiency

To calculate the protein cost of the HY pathway, the glucose uptake rate was fixed at 1 mmol/gDW/h, and the non-growth associated maintenance energy (NGAM) reaction was maximized. The total protein pool reaction was then minimized with fixing the NGAM reaction at the maximized value. The minimized flux through the total protein pool reaction is the protein cost of the HY pathway for converting one glucose to ATP. As for the protein cost calculation of LY pathway, glucose uptake rate was fixed at 1 mmol/gDW/h, the ethanol production was maximized. Then the ethanol exchange rate was fixed at the maximized value, and NGAM was maximized. After that, NGAM was also fixed at the maximized value, and total protein pool was minimized to calculate the protein cost for LY pathway. We also examined the flux distribution to ensure that other energy producing pathways are all inactive during this simulation. Protein efficiency is defined as the protein cost for producing one flux ATP in both pathways.

### Code and data availability

To facilitate further usage, we provide all codes, example and detailed instruction in GitHub repository: https://github.com/SysBioChalmers/DLKcat. Protein sequence fasta files, deep learning predicted *k*_cat_ values, classcial-ecGEMs, DL-ecGEMs and *Posterior*-mean-ecGEMs for 343 yeast/fungi species are available as Supplementary Dataset on the zenodo: https://doi.org/10.5281/zenodo.5164210.

## Author contribution

F.L, L.Y.,H.L. and J.N. designed the research. F.L. and L.Y. performed the research. F.L, L.Y., Y.C., G.L., E.K. and J.N. analyzed the data. L.Y. and M.E. collected the *k*_cat_ data. F.L, L.Y., H.L, G.L., Y.C., M.E., E.K. and J.N. wrote the paper. All authors approved the final paper.

## Acknowledgement

This project has received funding from the Novo Nordisk Foundation (grant no. NNF10CC1016517), the Knut and Alice Wallenberg Foundation, and the European Union’s Horizon 2020 research and innovation program with projects DD-DeCaF (grant no. 686070). The computations were enabled by resources provided by the Swedish National Infrastructure for Computing (SNIC) at Chalmers Centre for Computational Science and Engineering (C3SE) and High Performance Computing Center North (HPC2N), partially funded by the Swedish Research Council through grant agreement no. 2018-05973.

## Competing interests

The authors declare no competing interests.

## Notes

https://doi.org/10.5281/zenodo.5164210

https://github.com/SysBioChalmers/DLKcat

### Competing Interest Statement

The authors have declared no competing interest.

## Reference

1. Chen, Y. & Nielsen, J. Energy metabolism controls phenotypes by protein efficiency and allocation. Proc. Natl. Acad. Sci. U. S. A. 116, 17592–17597 (2019).

2. Sánchez, B. J. et al. Improving the phenotype predictions of a yeast genome-scale metabolic model by incorporating enzymatic constraints. Mol. Syst. Biol. 13, 935 (2017).

3. Klumpp, S., Scott, M., Pedersen, S. & Hwa, T. Molecular crowding limits translation and cell growth. Proc. Natl. Acad. Sci. U. S. A. 110, 16754–16759 (2013).

4. Schomburg, I. et al. The BRENDA enzyme information system–From a database to an expert system. J. Biotechnol. 261, 194–206 (2017).

5. Wittig, U., Rey, M., Weidemann, A., Kania, R. & Müller, W. SABIO-RK: an updated resource for manually curated biochemical reaction kinetics. Nucleic Acids Res. 46, D656–D660 (2018).

6. Bar-Even, A. et al. The moderately efficient enzyme: evolutionary and physicochemical trends shaping enzyme parameters. Biochemistry 50, 4402–4410 (2011).

7. Chen, Y. & Nielsen, J. Mathematical modelling of proteome constraints within metabolism. Curr. Opin. Syst. Biol. (2021).

8. Davidi, D. & Milo, R. Lessons on enzyme kinetics from quantitative proteomics. Curr. Opin. Biotechnol. 46, 81–89 (2017).

9. Heckmann, D. et al. Machine learning applied to enzyme turnover numbers reveals protein structural correlates and improves metabolic models. Nat. Commun. 9, 1–10 (2018).

10. Nilsson, A., Nielsen, J. & Palsson, B. O. Metabolic models of protein allocation call for the kinetome. Cell Syst. 5, 538–541 (2017).

11. Kitchin, J. R. Machine learning in catalysis. Nat. Catal. 1, 230–232 (2018).

12. Shrivastava, A. D. & Kell, D. B. FragNet, a Contrastive Learning-Based Transformer Model for Clustering, Interpreting, Visualizing, and Navigating Chemical Space. Molecules 26, (2021).

13. Zrimec, J. et al. Deep learning suggests that gene expression is encoded in all parts of a co-evolving interacting gene regulatory structure. Nat. Commun. 11, 6141 (2020).

14. Kroll, A., Heckmann, D. & Lercher, M. J. Prediction of Michaelis constants from structural features using deep learning. Preprint at https://doi.org/10.1101/2020.12.01.405928 (2020).

15. Ryu, J. Y., Kim, H. U. & Lee, S. Y. Deep learning enables high-quality and high-throughput prediction of enzyme commission numbers. Proc. Natl. Acad. Sci. 201821905 (2019).

16. Kanehisa, M., Furumichi, M., Tanabe, M., Sato, Y. & Morishima, K. KEGG: new perspectives on genomes, pathways, diseases and drugs. Nucleic Acids Res. 45, D353– D361 (2017).

17. Yep, A., Kenyon, G. L. & McLeish, M. J. Saturation mutagenesis of putative catalytic residues of benzoylformate decarboxylase provides a challenge to the accepted mechanism. Proc. Natl. Acad. Sci. U. S. A. 105, 5733–5738 (2008).

18. Lin, Y.-H. T., Huang, C. L. V., Ho, C., Shatsky, M. & Kirsch, J. F. A general method to predict the effect of single amino acid substitutions on enzyme catalytic activity. Preprint at https://doi.org/10.1101/236265 (2017).

19. Bahdanau, D., Cho, K. & Bengio, Y. Neural machine translation by jointly learning to align and translate. Preprint at https://arxiv.org/abs/1409.0473v7 (2014).

20. Erion, M. D. et al. Purine nucleoside phosphorylase. 1. Structure-function studies. Biochemistry 36, 11725–11734 (1997).

21. feiranl, hongzhonglu, Domenzain, I. & Yuan, L. SysBioChalmers/Yeast-Species-GEMs: Yeast-Species-GEM. (2021). data sets. zenodo https://doi:10.5281/zenodo.4568962

22. Nam, H. et al. Network context and selection in the evolution to enzyme specificity. Science 337, 1101–1104 (2012).

23. Kryazhimskiy, S. & Plotkin, J. B. The population genetics of dN/dS. PLoS Genet. 4, e1000304 (2008).

24. Ringe, D. & Petsko, G. A. Biochemistry. How enzymes work. Science 320, 1428–1429 (2008).

25. Li, G. et al. Bayesian genome scale modelling identifies thermal determinants of yeast metabolism. Nat. Commun. 12, 1–12 (2021).

26. Van Hoek, P. I. M., Van Dijken, J. P. & Pronk, J. T. Effect of specific growth rate on fermentative capacity of baker’s yeast. Appl. Environ. Microbiol. 64, 4226–4233 (1998).

27. Bekiaris, P. S. & Klamt, S. Automatic construction of metabolic models with enzyme constraints. BMC Bioinformatics 21, 19 (2020).

28. Pfeiffer, T. & Morley, A. An evolutionary perspective on the Crabtree effect. Front. Mol. Biosci. 1, 17 (2014).

29. de Alteriis, E., Cartenì, F., Parascandola, P., Serpa, J. & Mazzoleni, S. Revisiting the Crabtree/Warburg effect in a dynamic perspective: a fitness advantage against sugar-induced cell death. Cell Cycle 17, 688–701 (2018).

30. Ata, Ö. et al. A single Gal4-like transcription factor activates the Crabtree effect in *Komagataella phaffii*. Nat. Commun. 9, 1–10 (2018).

31. Kamrad, S. et al. Pyruvate kinase variant of fission yeast tunes carbon metabolism, cell regulation, growth and stress resistance. Mol. Syst. Biol. 16, e9270 (2020).

32. Krebs, H. A. Rate control of the tricarboxylic acid cycle. Adv. Enzyme Regul. 8, 335–353 (1970).

33. Christen, S. & Sauer, U. Intracellular characterization of aerobic glucose metabolism in seven yeast species by ^13^C flux analysis and metabolomics. FEMS Yeast Res. 11, 263–272 (2011).

34. Blank, L. M., Lehmbeck, F. & Sauer, U. Metabolic-flux and network analysis in fourteen hemiascomycetous yeasts. FEMS Yeast Res. 5, 545–558 (2005).

35. Chen, K. & Arnold, F. H. Engineering new catalytic activities in enzymes. Nat. Catal. 3, 203–213 (2020).

36. Markel, U. et al. Advances in ultrahigh-throughput screening for directed enzyme evolution. Chem. Soc. Rev. 49, 233–262 (2020).

37. Loeb, D. D. et al. Complete mutagenesis of the HIV-1 protease. Nature 340, 397–400 (1989).

38. Lee, J. & Goodey, N. M. Catalytic contributions from remote regions of enzyme structure. Chem. Rev. 111, 7595–7624 (2011).

39. Tong, H., Küken, A., Razaghi-Moghadam, Z. & Nikoloski, Z. Characterization of effects of genetic variants via genome-scale metabolic modelling. Cell. Mol. Life Sci. 78, 5123–5138 (2021).

40. Chen, Y., Li, F., Mao, J., Chen, Y. & Nielsen, J. Yeast optimizes metal utilization based on metabolic network and enzyme kinetics. Proc. Natl. Acad. Sci. 118, (2021).

41. Kim, S. et al. PubChem Substance and Compound databases. Nucleic Acids Res. 44, D1202–13 (2016).

42. Chen, F., Yuan, L., Ding, S., Tian, Y. & Hu, Q.-N. Data-driven rational biosynthesis design: from molecules to cell factories. Brief. Bioinform. 21, 1238–1248 (2020).

43. The UniProt Consortium. UniProt: the universal protein knowledgebase. Nucleic Acids Res. 45, D158–D169 (2017).

44. Tsubaki, M., Tomii, K. & Sese, J. Compound-protein interaction prediction with end-to-end learning of neural networks for graphs and sequences. Bioinformatics 35, 309–318 (2019).

45. LeCun, Y., Bengio, Y. & Hinton, G. Deep learning. Nature 521, 436–444 (2015).

46. Dong, Q.-W., Wang, X.-L. & Lin, L. Application of latent semantic analysis to protein remote homology detection. Bioinformatics 22, 285–290 (2006).

47. Moretti, S., Tran, V. D. T., Mehl, F., Ibberson, M. & Pagni, M. MetaNetX/MNXref: unified namespace for metabolites and biochemical reactions in the context of metabolic models. Nucleic Acids Res. 49, D570–D574 (2021).

48. Shen, X.-X. et al. Tempo and mode of genome evolution in the budding yeast subphylum. Cell 175, 1533–1545 (2018).

49. Yang, Z. PAML 4: phylogenetic analysis by maximum likelihood. Mol. Biol. Evol. 24, 1586–1591 (2007).

50. Domenzain, I. et al. Reconstruction of a catalogue of genome-scale metabolic models with enzymatic constraints using GECKO 2.0. Preprint at https://doi.org/10.1101/2021.03.05.433259 (2021).

51. Popovic, M. Thermodynamic properties of microorganisms: determination and analysis of enthalpy, entropy, and Gibbs free energy of biomass, cells and colonies of 32 microorganism species. Heliyon 5, e01950 (2019).

52. Yu, R. et al. Nitrogen limitation reveals large reserves in metabolic and translational capacities of yeast. Nat. Commun. 11, 1881 (2020).

53. Metzl-Raz, E. et al. Principles of cellular resource allocation revealed by condition-dependent proteome profiling. Elife 6, (2017).

