## Supplementary Figure for "Deep learning based *k*_cat_ prediction enables improved enzyme constrained model reconstruction"

**Supplementary Figures for Deep learning based $k_{cat}$ prediction enables improved enzyme constrained model reconstruction**

Feiran Li[1, #], Le Yuan[1, 2, #], Hongzhong Lu[1], Gang Li[1], Yu Chen[1], Martin K. M. Engqvist[1], Eduard J Kerkhoven[1, 2], Jens Nielsen[1, 3, *]

1 Department of Biology and Biological Engineering, Chalmers University of Technology, Kemivägen 10, SE-412 96 Gothenburg, Sweden

2 Novo Nordisk Foundation Center for Biosustainability, Chalmers University of Technology , Kemivägen 10, SE-412 96, Gothenburg, Sweden

3 BioInnovation Institute, Ole Måløes Vej 3, DK2200 Copenhagen N, Denmark

### These authors contributed equally to this work: Feiran Li, Le Yuan.

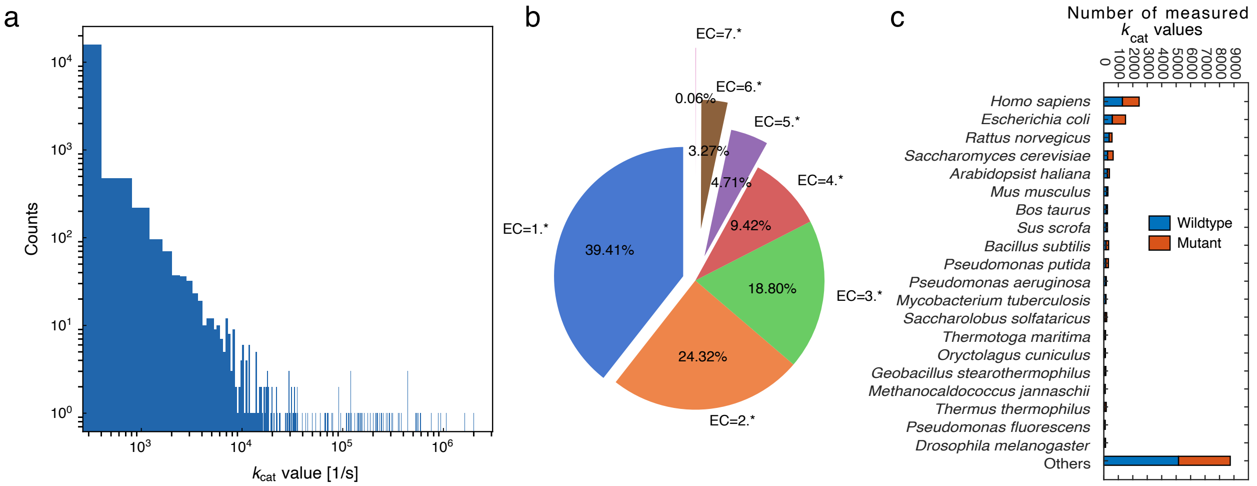


**Supplementary Figure 1** Data analysis for *in vitro* $k_{cat}$ values collected from the BRENDA and the SABIO-RK database after several rounds of data preprocessing, cleaning and combination. (a) Data distribution of *in vitro* $k_{cat}$ values. (b) Classification of *in vitro* $k_{cat}$ values based on the first digit of EC number. (c) Classification of *in vitro* $k_{cat}$ values based on species.

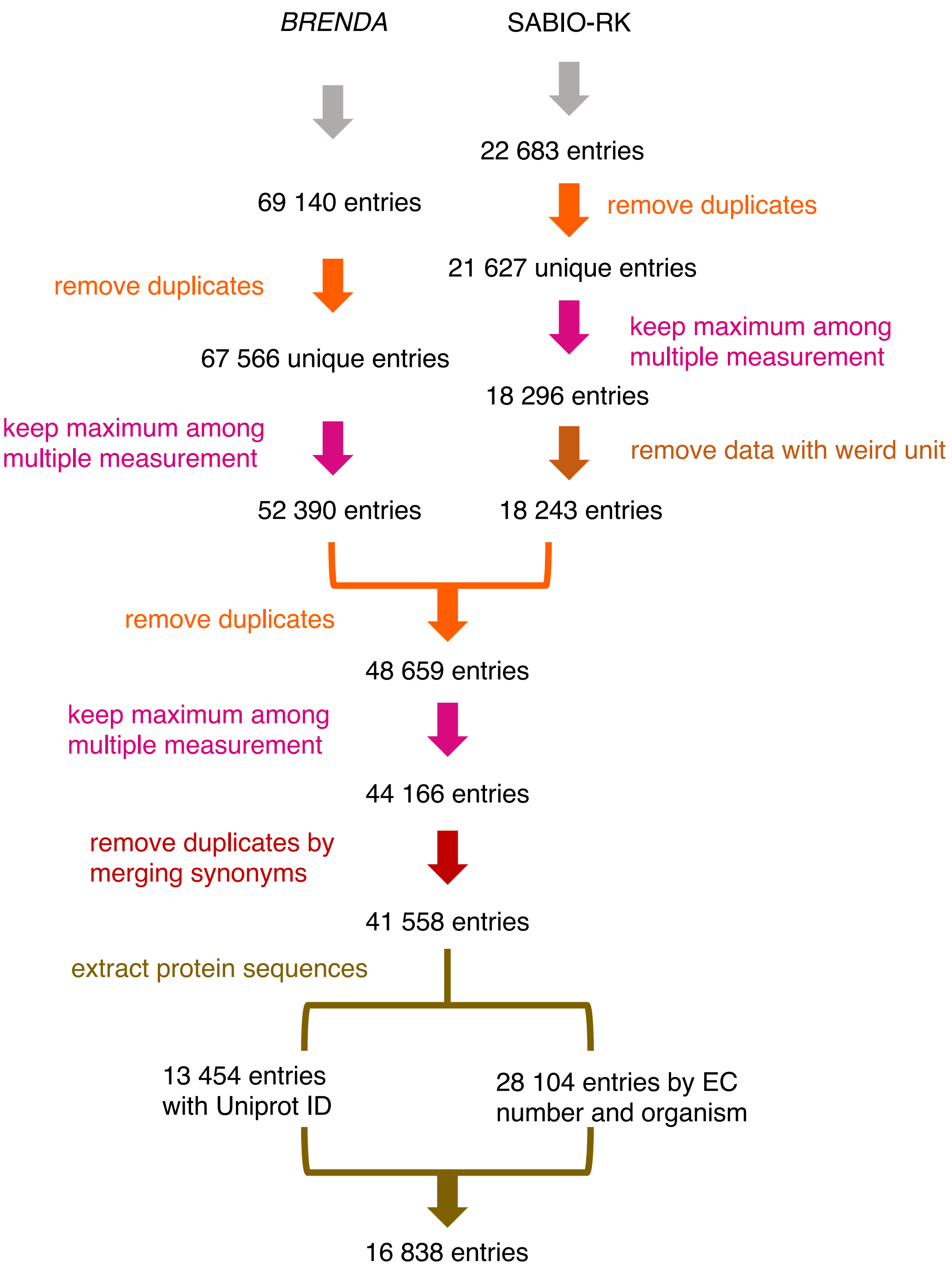


**Supplementary Figure 2** Data cleaning process for deep learning model input.

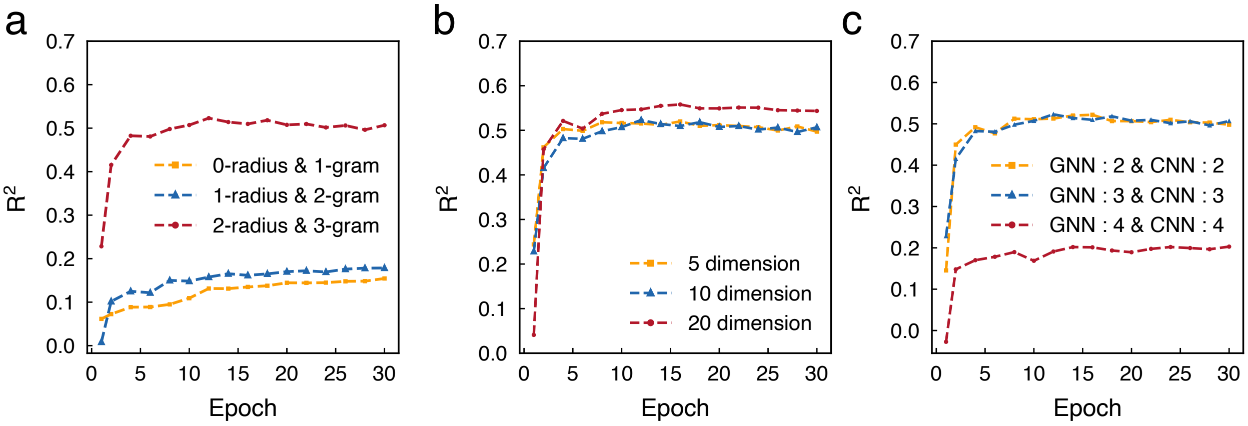


**Supplementary Figure 3** Learning curves with various hyperparameters on the validation dataset, including (a) various r-radius subgraphs and n-gram amino acids, (b) various dimension vectors and (c) various numbers of layers in GNN and CNN.

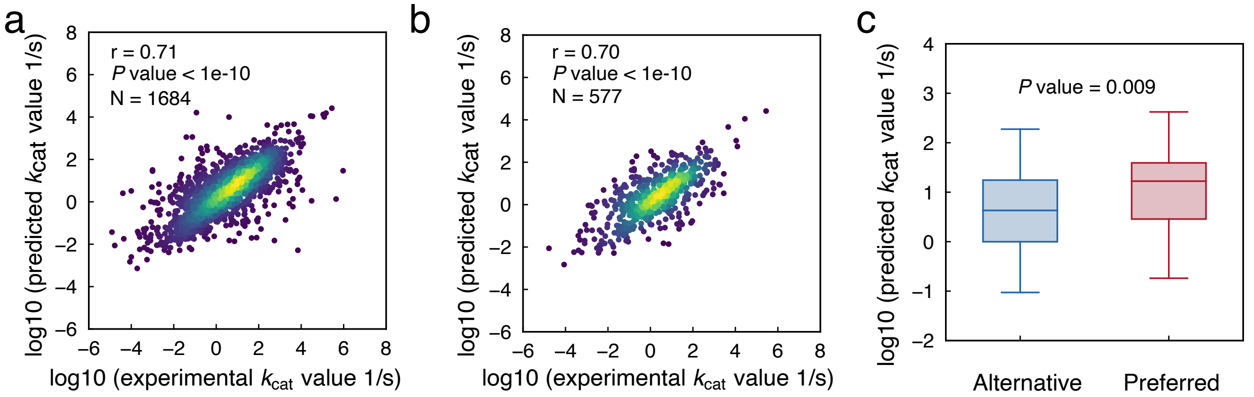


**Supplementary Figure 4** Deep learning model performance for $k_{cat}$ prediction on test dataset. (a) Performance of the final deep learning model on test dataset. The correlation between predicted $k_{cat}$ values and those present in the test dataset was evaluated. The brightness of color represents the density of data points. (b) Performance of the final deep learning model trained by GNN and CNN on subset of test dataset where the proteins sequence and the substrates were not involved in the training data set. (c) Enzyme promiscuity analysis on test dataset. For enzymes with multiple substrates, we divided the substrates as preferred and alternative by their experimental measured $k_{cat}$, and then plotted the predicted $k_{cat}$ values. A two-sided Wilcoxon rank sum test was used to calculate *P* value.

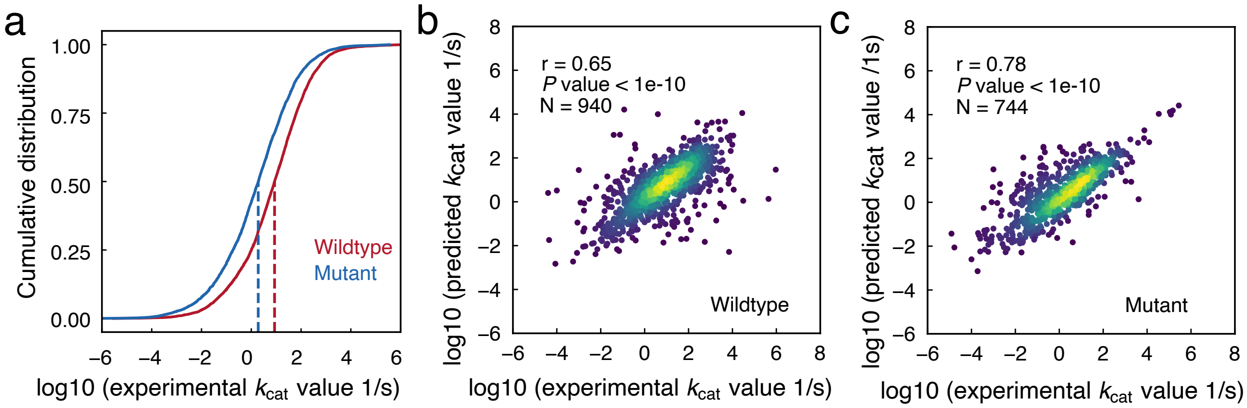


**Supplementary Figure 5** Comparison of $k_{cat}$ values for wildtype and mutated enzymes and prediction performance for both types of enzymes on test dataset. (a) Cumulative distribution of experimentally measured $k_{cat}$ values for wildtype and mutated enzymes. (b) Prediction performance of $k_{cat}$ values for all of the wildtype enzymes via deep learning model. The brightness of color represents the density of data points. (c) Prediction performance of $k_{cat}$ values for all of the mutated enzymes via deep learning model. The brightness of color represents the density of data points.

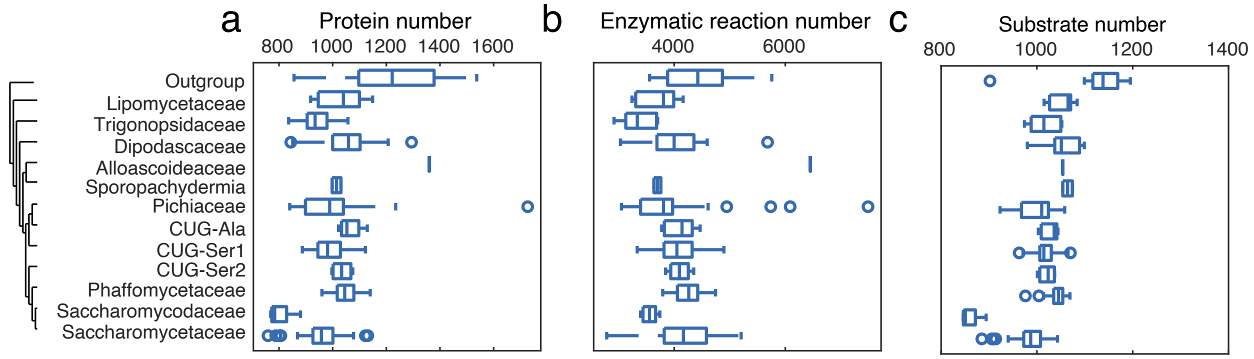


**Supplementary Figure 6** Comparison of (a) protein number, (b) enzymatic reaction number and (c) substrate number in deep learning predicted $k_{cat}$ profiles of 343 yeast/fungi GEMs. The x-axis represents the outgroup (11 species) together with 12 major clades divided by the genus-level phylogeny for 332 yeast species.

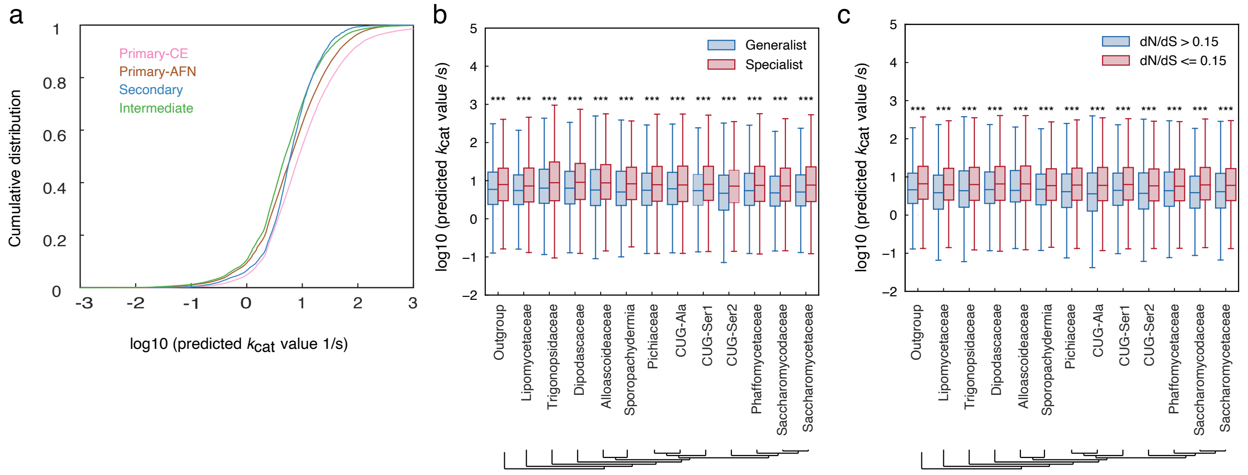


**Supplementary Figure 7** Evolution analysis of predicted $k_{cat}$ values for 343 yeast/fungi species. (a) Cumulative distribution of deep learning predicted $k_{cat}$ values for enzyme-substrate pairs in 343 yeast/fungi species belonging to different metabolic contexts. Abbreviations: CE, carbohydrate and energy; AFN, amino acids, fatty acids, and nucleotides. (b) Enzyme $k_{cat}$ values linked with generalist and specialist in the model for all 343 yeast/fungi species. (c) Enzyme $k_{cat}$ values linked with the ratio of non-synonymous to synonymous substitution (dN/dS) analysis for all of 343 yeast/fungi species. The x-axis represents the outgroup (11 species) together with 12 major clades divided by the genus-level phylogeny for 332 yeast species. The cutoff of 0.15 was set according to the distribution of dN/dS values in these species. *** means $P$ value < 0.001 in the correlation test analysis.

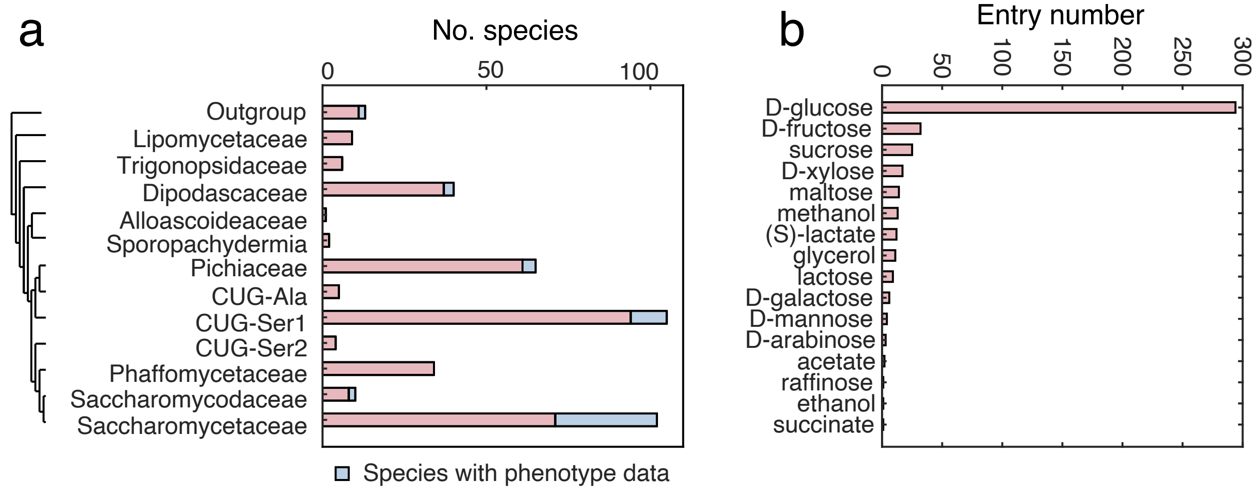


**Supplementary Figure 8** Bayesian approach related information. (a) Species analyzed in this study. (b) Collected growth rate data based on carbon source.

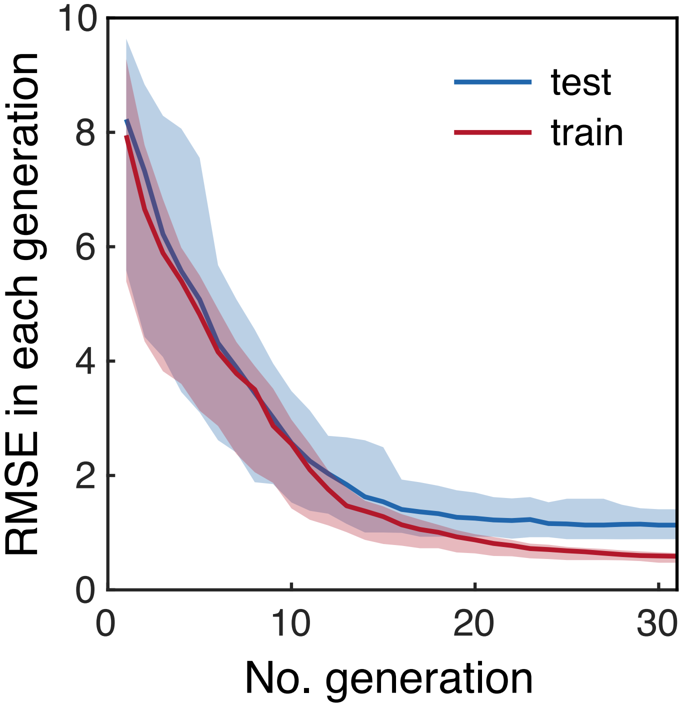


**Supplementary Figure 9** Validation of SMC-ABC approach. In this validation approach, data of 50% of experimental points were used to update the *Prior* and then tested on the remaining 50%. The split was done by first sorting all the data based on growth rates, then choosing the ones with even index for training and others for test. RMSE was calculated as described in Method. Lines indicate median values and shaded areas indicate regions between the 5-th and 95-th percentiles (n=100).

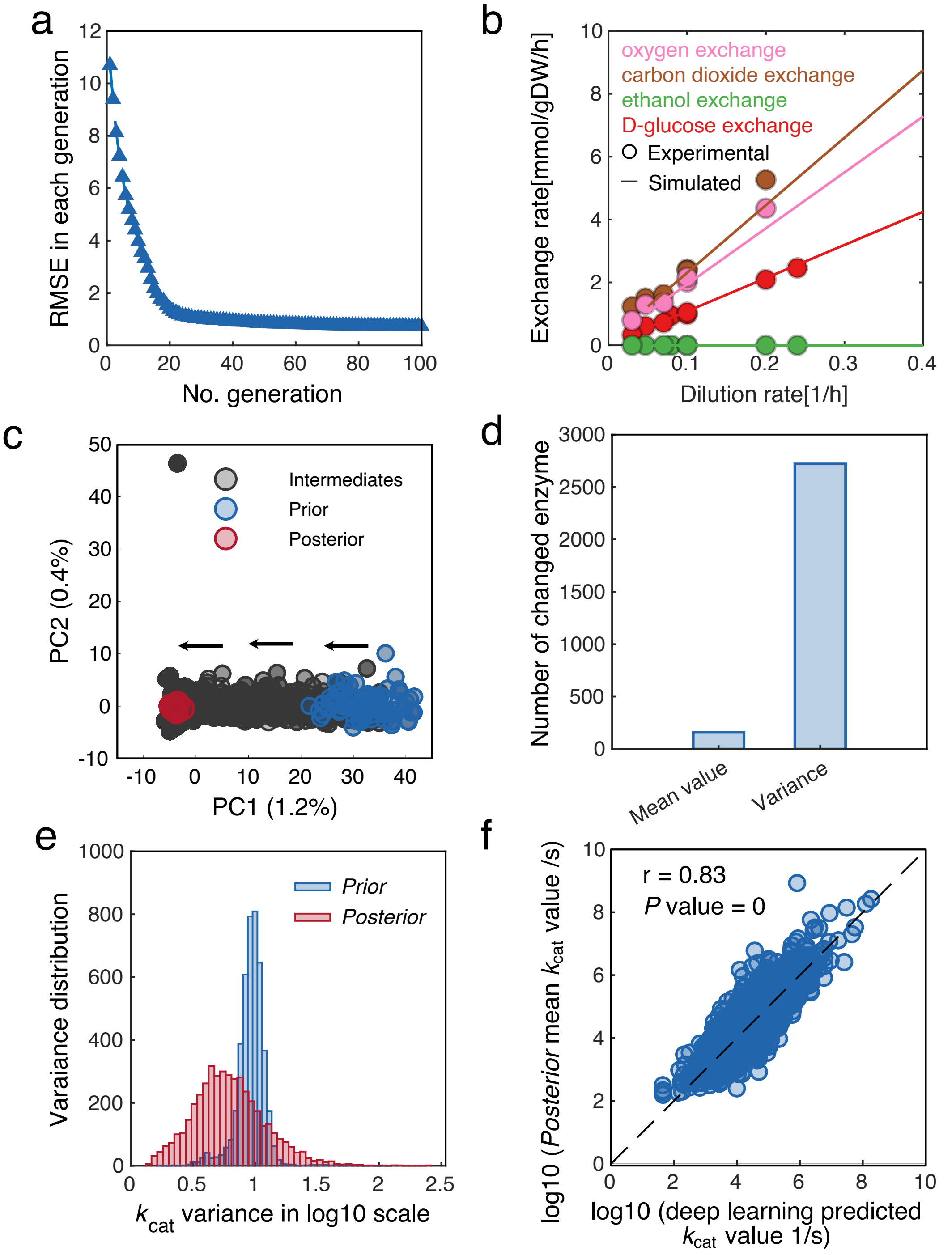
a
RMSE in each generation
No. generation
b
oxygen exchange
carbon dioxide exchange
ethanol exchange
D-glucose exchange
Experimental
Simulated
Exchange rate[mmol/gDW/h]
Dilution rate[1/h]
c
Intermediates
Prior
Posterior
PC2 (0.4%)
PC1 (1.2%)
d
Number of changed enzyme
Mean value
Variance
e
Prior
Posterior
Varaiance distribution
$k_{cat}$ variance in log10 scale
f
r = 0.83
P value = 0
log10 (Posterior mean $k_{cat}$ value /s)
log10 (deep learning predicted $k_{cat}$ value 1/s)

**Supplementary Figure 10** Bayesian modeling training performance for *Y. lipolytica* ecGEM. (a) The RMSE for phenotype of Bayesian training process. (b) Chemostat aerobic data and *Posterior*-mean-ecGEM simulations at increasing dilution rate. $k_{cat}$ values in the *Posterior*-mean-ecGEMs here is mean values from 100 sampled *Posterior* datasets after the Bayesian training process. (c) Principal component analysis (PCA) for $k_{cat}$ sets sampled in the Bayesian approach. Each parameter in the set was standardized by subtracting the mean and then be divided by the standard deviation before PCA. *Prior* datasets are highlighted in blue, while *Posterior* datasets are highlighted in red. All other datasets were termed as "intermediate" and marked in gray. (d) The number of enzymes with a significantly changed mean (Šidák adj. Welch's t test p value < 0.01, two-sided) and variance (Šidák adj. one-tailed F-test p value < 0.01) between sampled *Prior* and *Posterior* $k_{cat}$ datasets. (e) Variance distribution comparison for *Prior* and *Posterior* distribution. (f) Correlation analysis of deep learning predicted $k_{cat}$ values and *Posterior* $k_{cat}$ mean values.

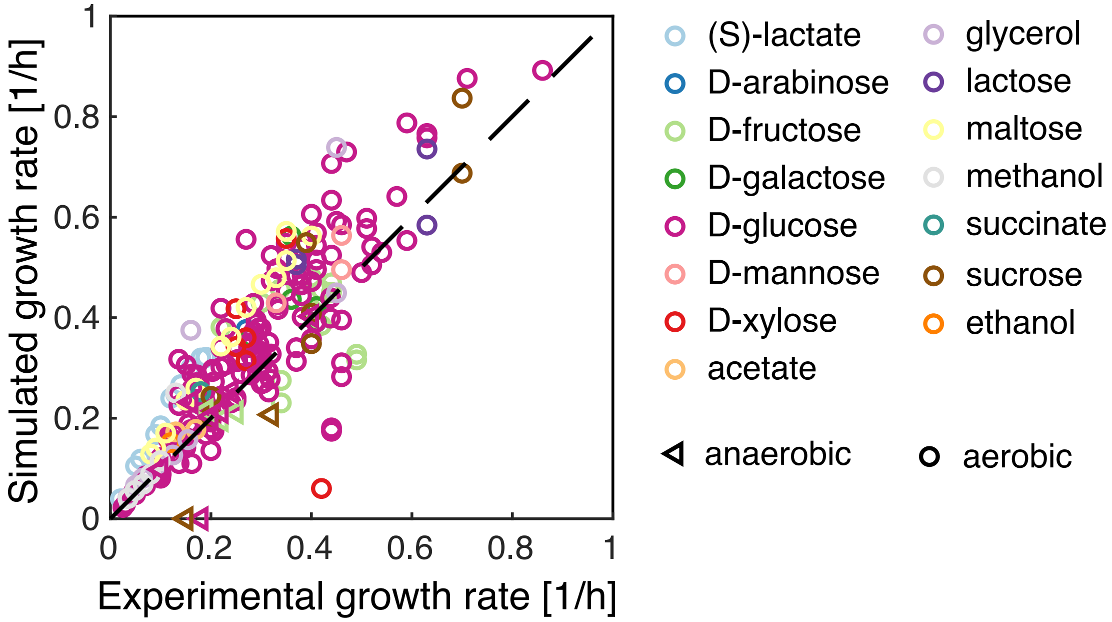


98

99 **Supplementary Figure 11** Growth prediction for *Posterior*-mean-ecGEMs on different carbon

100 sources. This figure is detailed version of Figure 5d.

101
